# Genome structural variants shape adaptive success of an invasive urban malaria vector *Anopheles stephensi*

**DOI:** 10.1101/2024.07.29.605641

**Authors:** Alejandra Samano, Naveen Kumar, Yi Liao, Farah Ishtiaq, Mahul Chakraborty

## Abstract

Global changes are associated with the emergence of several invasive species. However, the genomic determinants of the adaptive success of an invasive species in a new environment remain poorly understood. Genomic structural variants (SVs), consisting of copy number variants, play an important role in adaptation. SVs often cause large adaptive shifts in ecologically important traits, which makes SVs compelling candidates for driving rapid adaptations to environmental changes, which is critical to invasive success. To address this problem, we investigated the role SVs play in the adaptive success of *Anopheles stephensi*, a primary vector of urban malaria in South Asia and an invasive malaria vector in several South Asian islands and Africa. We collected whole genome sequencing data from 115 mosquitoes from invasive island populations and four locations from mainland India, an ancestral range for the species. We identified 2,988 duplication copy number variants and 16,038 deletions in these strains, with ∼50% overlapping genes. SVs are enriched in genomic regions with signatures of selective sweeps in the mainland and invasive island populations, implying a putative adaptive role of SVs. Nearly all high-frequency SVs, including the candidate adaptive variants, in the invasive island populations are present on the mainland, suggesting a major contribution of existing variation to the success of the island populations. Among the candidate adaptive SVs, three duplications involving toxin-resistance genes evolved, likely due to the widespread application of insecticides in India since the 1950s. We also identify two SVs associated with the adaptation of *An. stephensi* larvae to brackish water in the island and two coastal mainland populations, where the mutations likely originated. Our results suggest that existing SVs play a vital role in the evolutionary success of *An. stephensi* in new environmental conditions.

## Introduction

Invasive population expansion of a species often entails rapid adaptation to novel environments (Reznick and Ghalambor 2001). While ecological factors play a role in this process, increasing evidence supports the important contribution of genetic variation to invasive success (Lee 2002). Adaptation over a short evolutionary time relies on mutations conferring large fitness advantages in the new environments (Bomblies and Peichel 2022). Such mutations could arise *de novo* in the invasive population or through standing genetic variation in the source population (Prentis et al. 2008). Understanding the nature of these mutations and the relative role of novel and preexisting genetic variation to invasive range expansion is critical to predicting the potential for future invasion events. With the recurrent emergence of invasive species across the globe, it has become increasingly important to investigate the genetics of invasive success (Bohlen et al. 2004; Morrison et al. 2004; Hayes et al. 2008).

*Anopheles stephensi* is a primary vector of urban malaria in the Indian subcontinent and Middle East. However, the species is highly invasive and has spread rapidly to new islands, countries, and continents separated by natural barriers. For example, *An. stephensi* has invaded the Indian islands Lakshadweep and the country of Sri Lanka in the last 25 years (Sharma and Hamzakoya 2001; Dharmasiri et al. 2017). Recently, *An. stephensi* was also detected in Djibouti, a country in the Horn of Africa, in 2012 and has since been found in Ethiopia, Somalia, Sudan, and Nigeria (Carter et al. 2018; Kolaczinski et al. 2021; Abubakr et al. 2022; Hemming-Schroeder and Ahmed 2023). Unless controlled urgently, this invasive vector is predicted to spread all over Africa, invading most African countries and putting nearly 126 million people at risk (Takken and Lindsay 2019; Sinka et al. 2020). The threat notice of *An. stephensi* spread issued by the World Health Organization (WHO) in 2019 further underscores the seriousness of this situation.

*An. stephensi* has adapted to various anthropogenic changes and selective pressures in its native and invasive range in Asia and Africa, making it a formidable obstacle in controlling urban malaria. A primary concern is its resistance to diverse insecticides like DDT, malathion, dieldrin, and deltamethrin in nearly all populations, including South Asia, the Middle East, and Africa (Enayati et al. 2020). Another concern is its adaptation to breed in man-made habitats such as freshwater storage tanks and wells (Sinka et al. 2020). Additionally, *An. stephensi* have been found breeding in brackish water in tsunami-inundated coastal villages on the south coast of India and the recently invaded island country, Sri Lanka (Gunasekaran et al. 2005; Surendran et al. 2019). These adaptations have enabled the rapid range expansion further into urban areas on the Indian subcontinent and the surrounding islands (Dharmasiri et al. 2017). However, the genomic basis of these adaptations, which play an important role in the invasive spread of this species, remains unknown, impeding effective chemical, ecological, or genetic control strategies.

Genome structural variants (SVs) like duplication, deletion, transposition, and inversion of large (>100 bp) sequences provide a major source of adaptive genetic variation (Daborn et al. 2002; Van’t Hof et al. 2016; Chakraborty et al. 2018; Harringmeyer and Hoekstra 2022). Gene duplications play an important role in the evolution of insecticide resistance in various insect species (Anthony et al. 1998; Newcomb et al. 2005; Zimmer et al. 2018). Metabolic resistance to insecticides can occur through the amplification of detoxification genes such as cytochrome P450s, esterases, or glutathione S-transferases (Liu 2015). For example, duplication and transposable element (TE) insertions in the cytochrome P450 gene *Cyp6g1* are associated with increased resistance to DDT in *Drosophila melanogaster* (Daborn et al. 2002; Schmidt et al. 2010). Similarly, a cytochrome P450 gene *Cyp9M10* duplication is linked to metabolic resistance to permethrin in *Culex quinquefasciatus* (Hardstone et al. 2010; Wilding et al. 2012). Resistance can also result from mutations affecting the target sites of insecticides, such as the acetylcholinesterase gene *Ace-1* (Liu 2015). In both *Culex* and *Anopheles* mosquitoes, tandem duplications that pair a wildtype and resistant copy of *Ace-1* lead to resistance to carbamate and organophosphate insecticides (Labbé et al. 2007; Weetman et al. 2015).

An examination of the population genomics of SVs in *An. stephensi* in established and new populations can elucidate the contribution of SVs *to* adaptive evolution and range expansion in this species. However, we only know about a few inversion polymorphisms studied using polytene chromosomes in the South Asian and Middle Eastern populations (Coluzzi 1972; Mahmood and Sakai 1984). Thus, the contribution of SVs in the adaptive evolution of traits relevant to invasion success, such as insecticide resistance or tolerance to brackish water, remains unknown. To investigate the adaptive significance of SVs in *An. stephensi*, we analyzed whole genome sequence data from 115 individual mosquitoes from four mainland locations in India and an archipelago where *An. stephensi* recently invaded, similar to its invasion of Africa (Fig. 1A). Using a population genomic map of duplications, deletions, and TE insertions, we show that SVs provide a significant source of putative adaptive genetic variation in mainland and island populations. We further show that most SVs, including the candidate adaptive SVs, in the island populations came from the mainland. We highlight several adaptive copy number variants that are candidates for driving rapid adaptations in the ancestral and the new *An. stephensi* populations in India.

**Figure 1.**
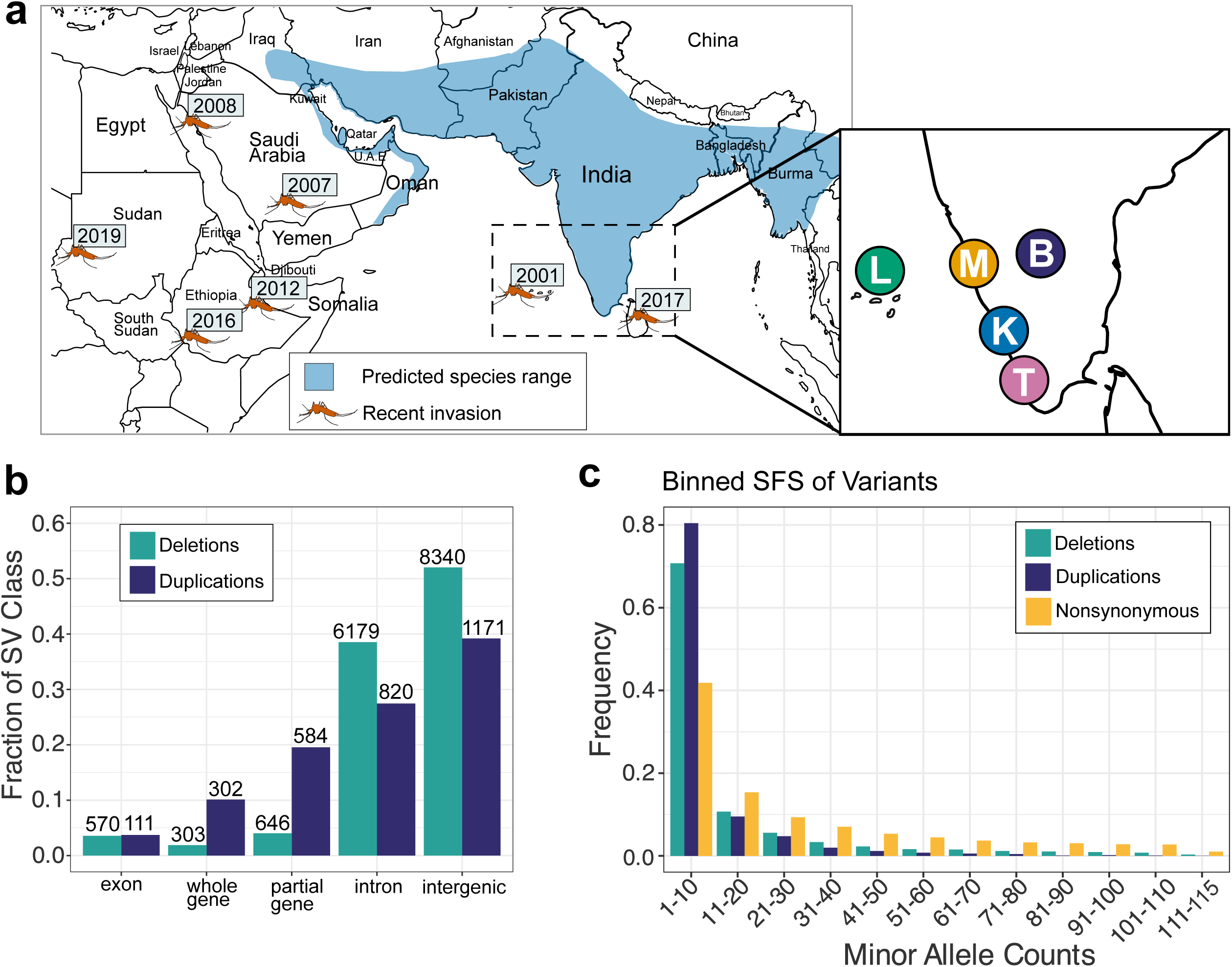
Structural variants in an island and four mainland populations of *An. stephensi*. **a**. Predicted distribution and recent invasive spread. Populations examined in this study: Bangalore (B), Mangalore(M), Kochi(K), Trivandrum(T), Lakshadweep(L). **b**. Distribution of duplication and deletion SVs in five genomic contexts: exonic (fully contained within an exon), whole gene (overlaps at least one complete gene), partial gene (overlaps gene but not completely), intronic (fully contained within an intron), intergenic (fully contained within an intergenic region). **c**. Binned minor allele counts of duplications, deletions, and nonsynonymous SNPs.

## Results

### Landscape of SVs in *An. stephensi* populations

To construct the genome-wide map of duplications and deletions, we mapped 150 bp paired-end Illumina reads (average coverage 20X assuming a genome size of 235 Mbp) from individual wild-caught *An. stephensi* mosquitoes to the *An. stephensi* reference genome (Chakraborty et al. 2021)(Supplementary Table 1). Short reads often miss duplications and report false positives (Chakraborty et al. 2018). Thus, we investigated the genotyping strategy to minimize false negative and false positive rates. We found that the SV detection strategy based on read pair orientation and split-read mapping implemented in the software Delly (Rausch et al. 2012) produced a reasonably reliable map of duplication and deletions for *D. melanogaster* genomes (Supplementary Fig. 1). We used that approach here for SV detection (see Methods for details). Additionally, using publicly available long and short reads for an *An. stephensi* strain, we validated 89% (40/45) duplicates and 91% (41/45) deletions from a randomly selected subset of SVs (see Methods). Thus, despite the limitations of short reads in SV detection (Chakraborty et al. 2018), the error rate in our SV dataset is low.

In the 115 samples, we discovered 2,988 duplications and 16,038 deletions with respect to the reference genome (Chakraborty et al. 2021), with an average of 196 and 2,105 duplications and deletions per individual (Supplementary Fig. 2). On average, duplications are longer than deletions (median length of duplications is 2,245 bp and 316 bp for deletions, Wilcoxon rank sum test, *p*-value < 2.2×10^-16^), similar to their counterparts in *D. melanogaster* (Emerson et al. 2008). Duplication CNVs are more prevalent on the X chromosome than on the autosomes (Proportion test, *p*-value < 2.12×10^-4^), which could be because the X chromosome has a lower effective population size, so selection is weaker, leaving more mutations. The distribution of SVs varies between genic and intergenic regions (Fig. 1B). We classified all SVs into five mutually exclusive groups: intronic, exonic, intergenic, whole gene, and partial gene (see Methods). Among the SVs, 3.2% overlapped complete genes, 46.8% involved partial genes, and the rest were in intergenic sequences. SVs involving partial genes and contained in exons are significantly depleted, which indicates that these are strongly deleterious and are eliminated from these regions by purifying selection (Fisher’s exact test, *p*-value < 2.2×10^-16^). We observed a smaller proportion of whole and partial gene deletions than duplications, consistent with the loss of a gene being more deleterious than copying it (Proportion test, *p*-value < 2.2×10^-16^).

About 5% (883/16,921) of the deletions completely overlapped annotated transposable element sequences in the reference genome, suggesting these deletions could represent polymorphic TE insertions in the reference genome (Supplementary Fig. 3-4). The abundance of polymorphic TE insertions in a single genome is similar to that of *D. melanogaster* (Cridland et al. 2013; Chakraborty et al. 2019), implying an extensive structural genetic variation in *An. stephensi* populations due to TE activities. Only 5 TE insertions are in exons, consistent with the harmful effects of TEs disrupting genic sequences. However, 32.16% (284/883) of the TE insertions are in introns, some of which could affect gene expression (Cridland et al. 2015). For example, a 203 bp long terminal repeat (LTR) retrotransposon fragment in the first intron of *Ace-2*, a sex-linked paralog of the insecticide resistance gene *Ace–1 (Hemingway and Ranson 2000)*, is segregating at high frequencies (54-89%) in all populations (Supplementary Fig. 5). The first intron of a gene is often enriched with regulatory sequences (Park et al. 2014). LTR TE fragments proximal to a gene can upregulate its expression (Daborn et al. 2002; Chakraborty et al. 2018). Thus, the LTR fragment might alter *Ace-2* gene expression.

### Natural selection on SVs

To understand the selective forces acting on the SVs, we compared the minor allele frequency spectra of duplications and deletions with that of nonsynonymous single nucleotide polymorphisms (nsSNPs). Nonsynonymous SNPs (nsSNPs) change proteins and are considered deleterious on average (Huber et al. 2017). The allele frequencies of SVs are skewed more towards low frequency than nsSNPs (*p*-value 2.2×10^-16^, *χ*^2^ test between frequency of SVs and nsSNPs), suggesting stronger purifying selection acting on the SVs on average than the nonsynonymous SNPs (Fig. 1C). However, many SVs (166 duplicates and 2,543 deletions) segregate at a high or intermediate frequency (>0.25) and could include candidates for mutations evolving under positive or balancing selection (Supplementary Fig. 6). Interestingly, we observed a two-fold enrichment of complete gene duplications in this subset of SVs compared to the genome-wide proportion (19.3% of >25% frequency vs. 10.1% of all duplicates, Fisher’s Exact Test, *p*-value 6.2×10^-4^). This enrichment could be due to the higher probability of full gene duplicates having a beneficial function than partial genes or noncoding intergenic sequences. Among the high-and intermediate-frequency full gene duplications, 25 involved protein-coding genes, and 7 encompassed long non-coding RNAs. Whole gene deletions are generally more harmful than complete gene duplicates. Consistent with this, we identified only eight complete gene deletions among SVs segregating at allele frequencies greater than 25%.

Allele frequency of a mutation could rise due to positive selection but also increase due to neutral evolutionary processes. A signature of a selective sweep (Nielsen et al. 2005; Stephan 2019) at or near a high-frequency SV can further support its adaptive significance. To investigate SV mutations that evolved under positive selection, we examined signatures of a selective sweep using composite likelihood ratio (CLR) (Nielsen et al. 2005; DeGiorgio et al. 2016) near SVs in each of the five populations (Supplementary Fig. 7). Genomic windows with high CLR values are likely to contain adaptive variants. Thus, we examined the abundance of high (>25%) frequency SVs at genomic windows with the top 5% genomewide CLR values. We found SVs segregating at 25% or above allele frequencies are enriched (*p*-values 1×10^-5^-0.045) at these 5-kb windows with high CLR values (Supplementary Fig. 8). Some CLR peaks exist in all populations, whereas others are specific to populations, likely indicative of local adaptations (Fig. 2). Notably, the value of the CLR statistic across the genome is lower for Lakshadweep than the mainland populations, which is likely due to the effects of population bottleneck and growth on CLR (Nielsen et al. 2005; Alachiotis and Pavlidis 2018). GO enrichment analysis of genes affected by SVs associated with CLR peaks reveals several overrepresented terms associated with insecticide resistance, such as oxidoreductase activity, heme binding, and tetrapyrrole binding (Supplementary Fig. 9). We also identified several high-frequency (>0.8) nsSNPs associated with CLR peaks. These SNPs are within genes implicated in insecticide resistance (*Cyp9f2*), olfaction (*Nrf-6*), and immune response (*CD81*)(Ahmed et al. 2021; Derilus et al. 2023; Heilig et al. 2024) (Supplementary Table 6).

**Figure 2.**
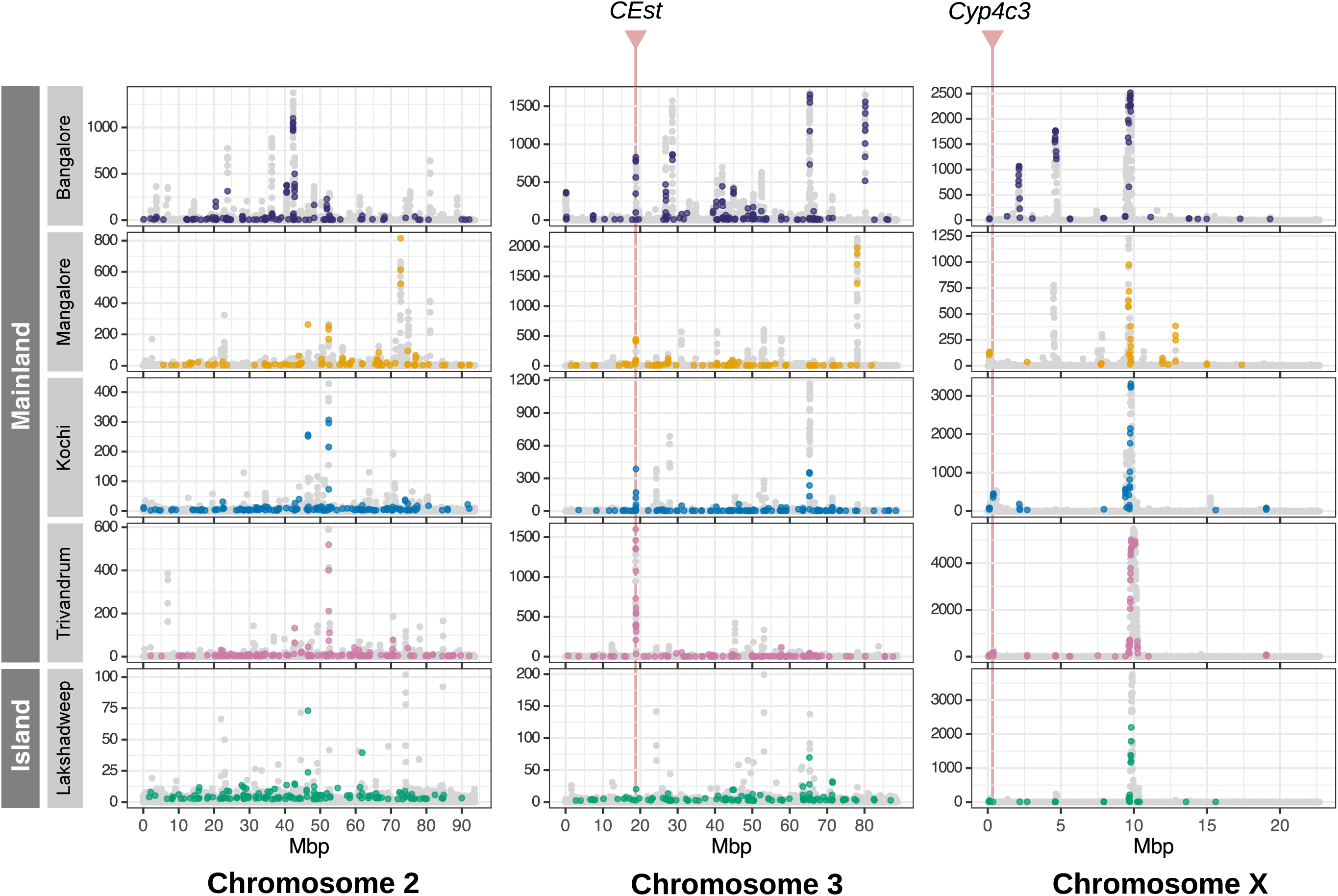
The distribution of composite likelihood ratio (CLR) statistic from the genome-wide scans for selective sweeps using SweepFinder2. Points are colored if the window CLR is in the 95th percentile and overlaps an SV with an allele frequency greater than 25% in that population. High-frequency duplications of carboxylesterases (CEst) and cytochrome P450 (*Cyp4c3*) genes are associated with CLR peaks in all populations.

### Adaptive evolution at insecticide resistance genes

Indian populations of *An. stephensi*, similar to other Asian and African populations, show widespread resistance to malathion and various pyrethroids (Enayati et al. 2020). Although the increased activity of beta carboxylesterase enzymes is thought to be responsible for this resistance, mutations underlying this enhanced detoxification activity are unknown (Ganesh et al. 2002; Prasad et al. 2017). One duplication SV we identified copies an 8.5 kb sequence on chromosome 3, completely overlapping two beta carboxylesterase genes (Fig. 3A). Based on the read-depth analysis, we estimated the duplicate allele harboring a staggering 15 copies of the genes, suggesting the origin of an insecticide resistance gene array (Chakraborty et al. 2021). This high-frequency duplicate allele overlaps a high CLR peak on Chromosome 3 in all populations, suggesting that the duplicate allele experienced a selective sweep in the recent past (Fig. 2). Reduced levels of nucleotide heterozygosity (π) and Tajima’s D around the duplicated region in the haplotype bearing the duplicate allele, but not the haplotypes carrying the reference allele, further supports the evolution of this duplicate under positive selection (Fig. 3A). The sequences near the duplicate allele are identical across populations, except for some rare SNPs, suggesting a single haplotype with the duplicate allele sweeping through the *An. stephensi* populations in India (Fig. 3C). The inferred selection coefficient (*s*) of the selective sweep based on the intensity of selection (ɑ) of the CLR peak is 0.15, suggesting a strong fitness advantage of this duplication. Despite strong selective advantage, the duplication is not fixed in any population, suggesting a recent origin of the mutation. Consistent with this prediction, an estimate of the age of the selective sweep (see Methods) based on the SNPs flanking the duplicate in the Trivandrum population suggests that the sweep likely occurred very recently, approximately 226.02 generations ago (188.62-722.74 generations, 95% confidence interval) (Supplementary Fig. 10). Amplification of esterases enhances organophosphate (OP) insecticide resistance in *Culex* and *Anopheles* mosquitoes (Hemingway and Karunaratne 1998). Thus, the duplication CNV we identified could explain the adaptive increase in carboxylesterase activity in malathion-and deltamethrin-resistant *An. stephensi* strains.

**Figure 3.**
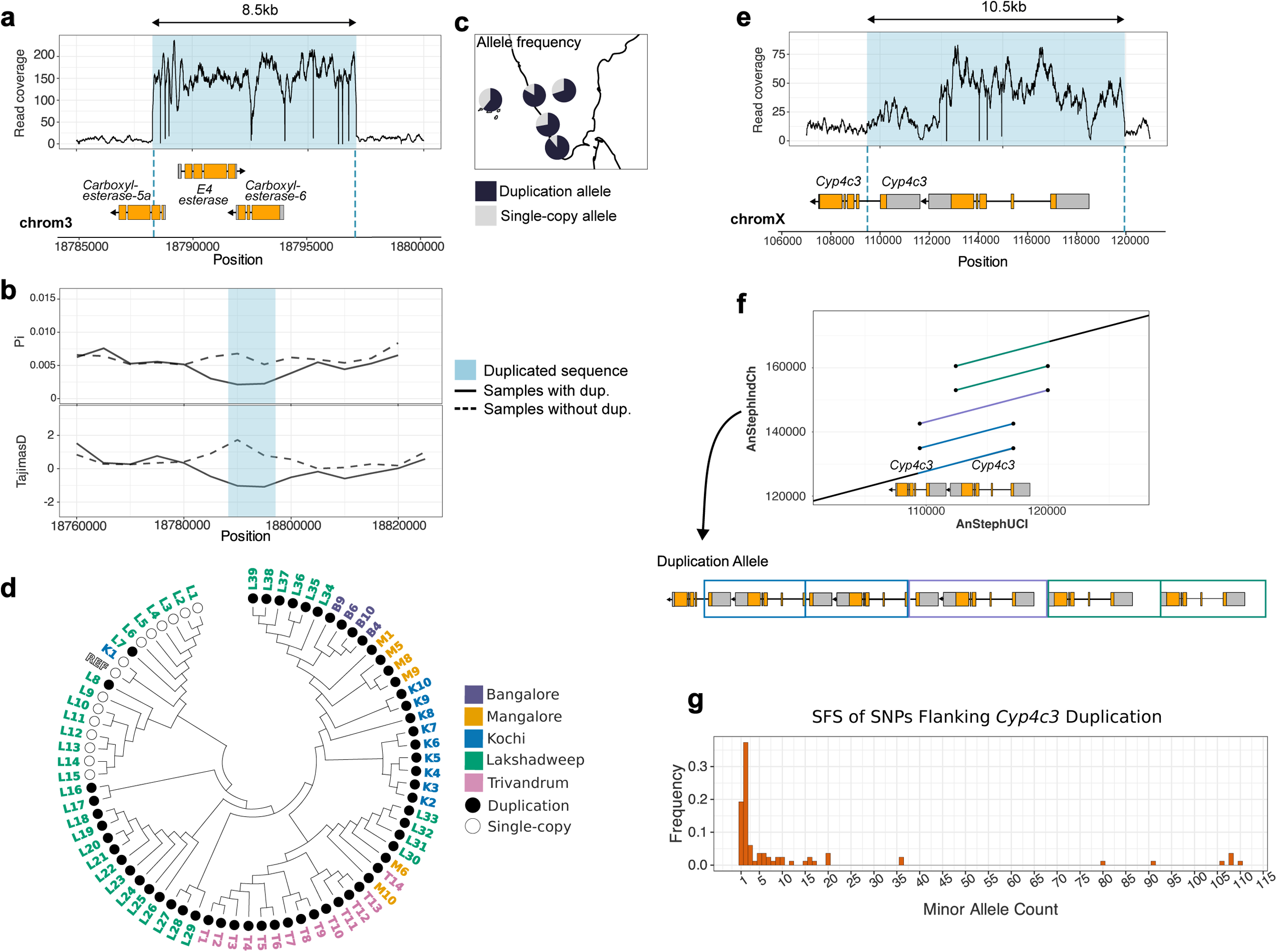
Candidate adaptive duplications for driving widespread insecticide resistance in *An. stephensi.* **a**. A duplication shown by the coverage of short reads mapped to a cluster of carboxylesterase (*CEst*) genes. **b**. Reduced nucleotide heterozygosity (π) and Tajima’s D flanking the duplication are indicative of recent positive selection. **c**. Allele frequency of *CEst* duplication allele in five populations. The duplication allele is high frequency in all populations. **d**. A gene tree constructed from the 20kb sequences flanking the *CEst* duplication. Only samples homozygous for the single-copy or duplicate allele are shown. All duplicate alleles except two form a single group consistent with a single origin of the duplicate allele. Two duplicate alleles clustering with the non-duplicate alleles likely represent recombination events. **e**. A duplication of two cytochrome P450 genes is fixed in all populations. **f**. A dot plot alignment between UCI reference assembly at *Cyp4c3* genes and duplication allele in the IndCh assembly. Lines represent an alignment between the genomes. Colored lines mark the copied sequences which are assembled to show the resulting gene structure of the duplicated region. **g**. A SFS of minor alleles for SNPs in the 20kb flanking regions of the *Cyp4c3* duplication.

We also identified a duplication CNV of a 10.5 kb sequence near the tip of the X chromosome, which is fixed in all populations and associated with a small CLR peak on the X chromosome (Fig. 2). The duplication copies two cytochrome P450 genes, both of which are similar to *D. melanogaster Cyp4C3*, expressed in the hindgut of feeding larvae (Chung et al. 2009). The low CLR value could be due to an old sweep, which allowed recombination to erode the effect of sweep on neutral variation linked to the adaptive variant. Due to the proximity of the duplication to the end of the X chromosome, very few SNPs are on the 5’ end of the duplication. However, an absence of intermediate frequency SNPs on the 5’ and 3’ ends (Fig. 3F) suggests, most likely, a single haplotype carrying the *Cyp4C3* duplication reached fixation in all *An. stephensi* populations. A more than two-fold read coverage indicates the presence of multiple copies of the genes in the duplicate allele. However, uneven coverage across the duplicate suggests different lengths of the individual copies within the duplicate (Fig. 3D). To examine the structure of the duplicate allele, we inspected the *Cyp4C3* gene in the published genome assembly of an *An. stephensi* strain (IndCh) collected from the Southern Indian city of Chennai in 2016 (Thakare et al. 2022). IndCh carries the fixed duplicate *Cyp4C3* allele, with six copies of the gene in duplicate (Fig. 3E). The *An. gambiae* ortholog of *An. stephensi Cyp4C3*, *Cyp4C26*, is overexpressed in pyrethroid-resistant *An. gambiae* strains in Kenya (Bonizzoni et al. 2015), suggesting a role of the *Cyp4C3* duplicate in the reduced susceptibility to pyrethroids like deltamethrin in *An. stephensi* (Tiwari et al. 2010).

Unlike the above two examples of gene duplications, for which a single allele most likely swept through all populations, we identified high-frequency gene duplication CNVs in a cluster of epsilon glutathione-S transferases (GSTe), for which two alleles are present in our samples. One allele shares the same breakpoints as a 3.6 kb tandem duplication reported in a laboratory-selected DDT-resistant strain of *An. stephensi* collected in India (Dykes et al. 2022). The duplicate copies two full-length GSTes and is associated with GST overexpression. We uncovered a second allele comprising a 2.9 kb duplication in the same cluster of genes, which also copies two complete GSTes. Interestingly, the duplicate reported by Dykes et al. segregates at high frequencies in Bangalore and is absent in Kochi. In contrast, the duplicate we uncovered segregates at high frequencies in Kochi and is lacking in Bangalore. In Mangalore, Trivandrum, and Lakshadweep, both duplications segregate at intermediate to high frequencies (3.6kb duplicate 17-22%, 2.9kb duplicate 22-53%). The read coverage pattern for 21 samples suggests they are heterozygotes of the two duplicate alleles or carry a recombinant allele. Thus, both GST duplication alleles could be linked to the DDT resistance observed in Asian and African populations of *An*. *stephensi (Raghavendra et al. 2017; Yared et al. 2020)*.

### Shared SV polymorphism in island and mainland populations

*An. stephensi* invaded Lakshadweep in the last 25 years, and insecticide resistance and its ability to adapt to a range of larval habitats likely helped the species colonize this island rapidly (Surendran et al. 2019; Ishtiaq et al. 2021). Thus, understanding the source of SVs in the Lakshadweep populations, including the variants contributing to the adaptive traits, can elucidate the relative contribution of segregating and *de novo* SV mutations in the rapid adaptations of *An. stephensi* in new habitats. In particular, we examined the proportion of SVs in island populations derived from mainland populations. While the specific adaptive mutations in these islands are unknown, such mutations often segregate at intermediate or high (>0.25) frequencies. Nearly all SV mutations in Lakshadweep segregating at frequencies 0.25 or above are also present in at least one mainland population (Fig. 4A, B). Approximately 25% of the SVs segregating at a frequency below 0.25 in the island population are not detected in the mainland populations, suggesting they are private to the island population or too rare to be detected in the mainland populations. The SV allele frequencies of the island population show a strong correlation with the allele frequencies of SVs from the coastal populations (Kochi & Trivandrum) but not with the SV allele frequencies of the inland (Bangalore & Mangalore) populations (Figure 4C, D). This pattern could be due to gene flow between the coastal and island populations, similar selective pressures in the two locations, or both.

**Figure 4.**
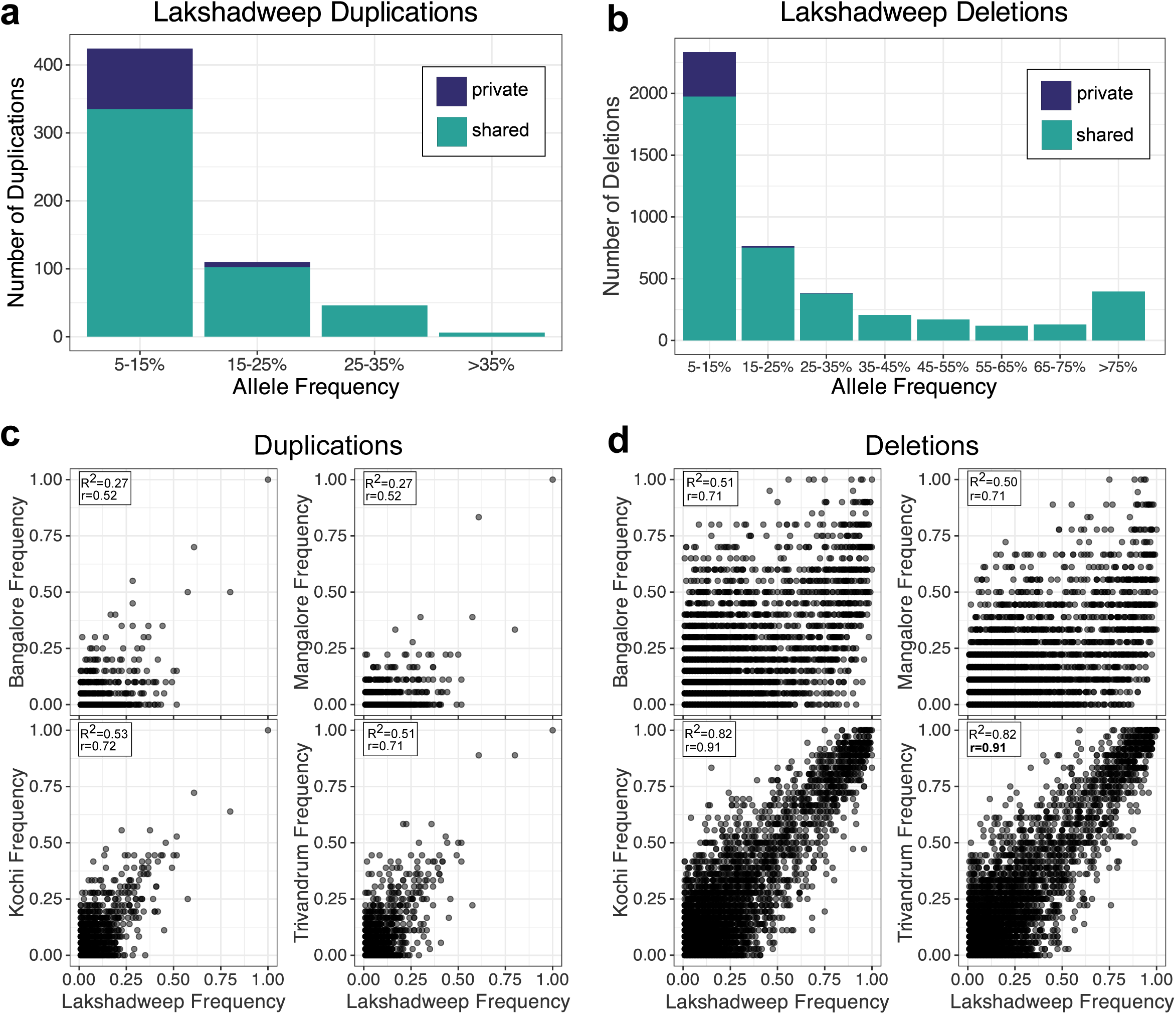
**a**. Allele frequency distribution of duplication SVs segregating at allele frequencies greater than 5% in the invasive island population. Duplications that are private or found only on the island segregate at low frequencies, while intermediate to high-frequency SVs are shared with at least one mainland population. **b.** Allele frequency distribution of deletion SVs segregating at allele frequencies greater than 5% in the invasive island population. As with duplications, deletions that are private to the island segregate at low frequencies. **c**. Allele frequency of duplications present in the island population is plotted against their frequency in each mainland population. **d**. Allele frequency of deletions present in the island population is plotted against their frequency in each mainland population. R^2^ and r represent the coefficient of determination and Pearson correlation coefficient, respectively, for the allele frequencies between the island and the mainland populations.

Consistent with the mainland being the source of putative adaptive SVs in the island, the esterase, *Cyp4c3, and GSTe* duplicates segregate at high frequencies in Lakshadweep islands (Fig. 3B). *An. stephensi* populations in coastal regions and recently invaded island populations often show tolerance to brackish water, although mutations responsible for these adaptations remain unknown. We found a 5.5 kb duplication on chromosome 3, which overlaps two larval cuticle protein A2B-like genes and a 1.5kb intronic deletion in a V-type proton ATPase gene (Supplementary Fig. 12A, B). Both mutations are present in the coastal populations from Kochi and Trivandrum, where they likely originated. Additionally, the cuticle protein and ATPase genes have been linked to salinity tolerance in *Anopheles* mosquitoes (Smith et al. 2008; Ramasamy et al. 2021; Sivabalakrishnan et al. 2023). Thus, these SVs could alter gene expression of the respective genes and contribute to *An. stephensi* adaptation to brackish water.

## Discussion

Using genome sequencing data from native and invasive populations of the urban malaria vector, *An. stephensi*, we have examined the role of SVs in adaptation to novel environments. We find that SVs present in mainland populations are the predominant source of high and intermediate-frequency SVs in invasive island populations. Although the SV allele frequency of island populations is driven by a combination of natural selection, ecological connectivity, demography, and gene flow, a high proportion of shared SV polymorphism for the intermediate or high-frequency category suggests that most, if not all, putatively adaptive SVs within this allele frequency category in island populations were derived from the mainland populations. In addition to high-frequency SVs, we observed low-frequency SVs in genes linked to insecticide resistance, host-seeking, mating behavior, and microbial resistance (Supplementary Tables 3,4,5). These SVs could be a potential source of genetic variation for contemporary or future adaptations.

Our results suggest SVs play an important role in the recurrent and rapid evolution of resistance to various insecticides in *An. stephensi* populations. The candidate duplicates, esterases, *Cyp4c3,* and *GST* likely evolved in the last 70 years due to the extensive use of insecticides in India after the Second World War. However, whether the timeline of the respective selective sweeps overlapped is unknown. The spread of the esterase duplicate could be driven by malathion, an OP insecticide that was introduced to India in 1969 and continues to be used in indoor residual sprays and outdoor fogging (Batra et al. 2005; Tiwari et al. 2010; Rahi and Sharma 2022). *An. stephensi* has been reported resistant to malathion in six states in India, including Karnataka, where two populations in this study were collected (Raghavendra et al. 2017). However, another possibility is that the spread of this duplication was driven by the more recent introduction of pyrethroid insecticides like deltamethrin. Although pyrethroid insecticide-treated mosquito nets (ITMN) were introduced in India under the National Anti Malaria Programme (NAMP) in the 1990s (Batra et al. 2005), pyrethroids continue to be used throughout India in ITMNs, indoor residual sprays, and outdoor fogging (Tiwari et al. 2010; Rahi and Sharma 2022). Given our estimation of a very recent selective sweep, it is likely that this duplication spread more recently as an adaptation to the extensive use of pyrethroids and could also explain the cross-resistance to malathion and pyrethroids reported in *An. stephensi*. The duplication overlapping the *Cyp4c3* genes could have been fixed due to the widespread use of synthetic pyrethroid insecticides for malaria vector control. These existing insecticide-resistant alleles from mainland populations helped the spreading of *An. stephensi* in Lakshadweep, where insecticides have been used to control this species. Invasive populations of *An stephensi* in Africa and Sri Lanka have already been found to be resistant to multiple classes of insecticides, suggesting that cross-resistance is a major contributor to the ongoing invasive success of the species. The GSTe duplications, similar to the DDT-resistance *Cyp6g1* alleles in *D. melanogaster*, also show how multiple SV alleles can potentially underlie the evolution of resistance to the same insecticide (Schmidt et al. 2010). Notably, the esterase, cytochrome P450 and GSTe gene duplications also show how tandem arrays of functional genes (Shukla et al. 2024) can arise under positive selection.

We also uncovered SVs that may contribute to local adaptation to the environmental conditions in the invasive and coastal populations. A duplication overlapping cuticle protein genes likely contributes to the larval adaptation to brackish water, which has been observed in *An. stephensi*, as well as other typically freshwater mosquito species. Salinity-tolerant forms of *Ae. aegypti* show increased expression of cuticle proteins, structural changes in larval cuticles, and increased resistance to temephos, an OP larvicide (Ramasamy et al. 2021; Sivabalakrishnan et al. 2023). This duplication may contribute to salinity tolerance in *An. stephensi*, though its role in larvicide resistance is unclear as larval resistance to temephos has not been confirmed in these populations. The deletion in the V-type proton ATPase gene may also contribute to salinity tolerance because changes in the localization of V-type proton ATPase and K+/Na+ ATPase play a role in osmotic regulation in saline-tolerant Anophelines (Smith et al. 2008). The rapid evolution of V-type proton ATPases in the copepod *Eurytema affinis* has been linked to its ability to invade freshwater habitats (Lee et al. 2011), which suggests that changes in this enzyme may contribute to adaptation to different salinities in *An. stephensi*.

Our results show SVs consisting of duplication, deletion, and TE insertions comprise a large proportion of uncharacterized functional genetic variation in *An. stephensi*. Importantly, we show evidence that putative adaptive variants from mainland populations segregate at high frequency in the newly colonized island population. This suggests *An. stephensi* populations invading new geographic locations will benefit from existing adaptive SVs and need not wait for new mutations to adapt to new environmental challenges, which helps *An. stephensi* to spread to new locations rapidly. Resistance to various insecticides and tolerance to diverse osmotic conditions are key adaptations in *An. stephensi* invasive populations in both Africa and Asia. Our results suggest SVs may play an important role in those adaptations. The ability to adapt to new environments is a crucial property of invasive species. Our results from *An. stephensi* suggests SVs can be a potential contributor to that property. However, our SV data based on short reads reveals only a fraction of total SVs in *An. stephensi* (Supplementary Fig. 1), potentially masking some adaptive variants (Chakraborty et al. 2018). Further research into the functional and fitness role of SVs using long reads in *An. stephensi* and other invasive species will help us better understand the role of SVs in invasion success.

## Materials & Methods

### Genomic data collection

Individual wild-type male and female mosquitoes were collected from four mainland populations and a population from the Lakshadweep archipelago in India (Supplementary Table 1) (Bangalore n = 10, Mangalore n = 9, Lakshadweep n = 60, Kochi n = 18, Trivandrum n = 18). The genomic DNA was isolated from homogenized whole mosquitoes using Qiagen Blood and Tissue kit (Qiagen). Data for the Bangalore and Mangalore strains are from a previously described sample (Thakare et al. 2022). All sequence data was generated using paired-end sequencing performed on an Illumina HiSeq 2500 platform (Illumina, San Diego, CA) at Tata Institute of Genetics and Society.

### Evaluation of SV Detection Strategies

We performed a benchmark analysis of SV detection methods to evaluate the performance of four short-read SV callers: Delly2 (Rausch et al. 2012), Lumpy (Layer et al. 2014), CNVnator (Abyzov et al. 2011), and Manta (Chen et al. 2016). Paired-end Illumina sequencing data (Shukla et al. 2024) from two inbred *Drosophila melanogaster* lines, A3 and A4, were sampled to 10, 15, and 25X coverage depth and aligned to the ISO1 reference genome (Hoskins et al. 2015). To simulate a heterozygous individual, we sampled the same three coverage depths but combined reads from A3 and A4 at equal proportions. We then applied the four short-read SV callers to identify duplications and deletions. The control dataset was generated using an assembly-based SV caller by comparing the highly contiguous genome assemblies of A3 and A4 to the reference ISO1 assembly(Chakraborty et al. 2019). The SVs identified with simulated short-read sequencing data at different depths were then compared with control data to evaluate the completeness (false negative rate) and accuracy (false positive rate) of each SV caller.

### SV genotyping

We used Trimmomatic v0.36 (Bolger et al. 2014) to remove adapters and trim low-quality regions from reads, followed by FASTQC v0.11.8 (Andrews 2010) to check read quality. Trimmed reads were then mapped to the AnSteph UCI reference genome (Chakraborty et al. 2021) using bwa-mem v0.7.17 with default parameters (Li 2013). Optical duplicates were marked and filtered out using Picard v2.23.9. To identify duplication and deletion SVs segregating in the five populations, we used Delly to implement read-pair orientation analysis (Rausch et al. 2012). We filtered SV calls, keeping only deletion and duplication variants between 100bp and 100kbp and excluding those with ‘LowQual’ flags using BCFtools v1.14 (Danecek et al. 2021). To merge SV calls across all 115 samples, we used Jasmine v1.1.5, which uses an SV proximity graph to merge variants present in multiple samples (Kirsche et al. 2023). We excluded SV calls with over 90% overlap with reference TEs. Deletion calls with over 90% reciprocal overlap with annotated TEs were considered polymorphic TE insertions and separated from the other deletion calls. The allele frequency of SVs was calculated from the merged call set using VCFtools v0.1.16 (Danecek et al. 2011).

Genotypes for candidate SVs were further inspected to confirm allele frequency and copy number. Copy number for the carboxylesterase duplication was determined from normalized read depth (RD) calculated by CNVnator v0.4.1, a coverage-based SV caller (Abyzov et al. 2011). The *Cyp4c3* duplication is reported as two separate, overlapping duplication calls, likely due to the different lengths of duplicated sequences within this mutation. 101/115 individuals are genotyped heterozygous for both duplications. However, visual inspection of this region of the BAM files in IGV (Thorvaldsdóttir et al. 2013) indicated that all samples show the coverage pattern observed in Figure 3D. Most of the samples incorrectly genotyped were male and, therefore, have lower X chromosome coverage, which could explain why this duplication was not called for these individuals. We also found several silent mutations within the duplicated sequence and in the flanking regions fixed in all populations, which we used as additional evidence to infer the fixation of this duplication allele in all populations.

### Validation

To evaluate the accuracy of our genotyping methods with *An. stephensi* data, we applied our SV detection pipeline to 20X Illumina sequencing data from IndCh (Thakare et al. 2022). We randomly selected 45 duplications (15 per chromosome) and 45 deletion SV calls found in both IndCh and our population SV dataset for further inspection. In particular, we checked whether long read and short read coverage for reads mapped to the UCI reference (Chakraborty et al. 2021) agreed with the filtered Delly calls, which use read-pair orientation and split-read mapping to infer the SVs. We found that 89% of duplications (40/45) and 91% of deletions (41/45) were supported by long and/or short read coverage (Supplementary Table 2). Some SVs supported by long reads are not supported by short reads and vice versa. This is because the IndCh strain is segregating for multiple haplotypes, and the DNA source for the short and long reads are separate, causing some haplotypes to be represented only in the long or short reads.

### SV ancestral state determination

To confirm that a variant corresponds to the derived state, a closely related outgroup species genome is needed to infer the ancestral state. The closest related species to *An. stephensi* with a high-quality genome assembly is *An. gambiae (Habtewold et al. 2023)*. We performed pairwise alignment between the *An. stephensi* and *An. gambiae* reference genomes (Chakraborty et al. 2021; Habtewold et al. 2023) using LASTZ v1.04.15 (Schwartz et al. 2003) and identified syntenic blocks using the UCSC Chain/Net pipeline (Kent et al. 2003). 50.44% of the *An. stephensi* genome aligned to *An. gambiae,* but only 5.1% of duplications and 13% of deletions in our callset fall within these syntenic regions. For this reason, we focused on determining the ancestral state of genic SVs, as coding regions are likely to be conserved between the two species, specifically genic SVs segregating at allele frequencies over 25% in a population. We used exonerate v2.4.0 (Slater and Birney 2005) to find ungapped alignments between the protein-coding sequence of completely duplicated or deleted genes and the *An. gambiae* genome. Additionally, we visually inspected syntenic regions using the JBrowse Genome Browser in Vectorbase. For gene duplications, if the gene was present in higher copies in *An. gambiae* than the *An. stephensi* reference, the duplication is inferred to be the ancestral state. For gene deletions, if the gene is present in fewer copies in *An. gambiae*, the deletion is inferred to be the ancestral state (a duplication in the *An. stephensi* reference).

### SNP genotyping

SNPs were identified using FreeBayes v1.3.5 (Garrison and Marth 2012) with the-C flag set to 2, specifying that a minimum of two reads must support the alternate allele to be called and the-0 flag, which excludes partially mapped reads from variant calling. Indels and low-quality variants were filtered out, and the remaining SNPs were annotated to predict the functional effect using SnpEff (Cingolani et al. 2012).

### Detection of Selective Sweeps

To detect genomic regions with signatures of recent positive selection, we used SweepFinder2, which performs genome-wide scans for selective sweeps by comparing the observed patterns of genetic diversity to a null model generated from the genome-wide frequency spectrum (Nielsen et al. 2005; DeGiorgio et al. 2016). The composite likelihood test implemented by SweepFinder2 has been found to have high power in detecting recent selective sweeps while being robust to demographic factors (Nielsen et al. 2005). Using the quality-filtered SNP data, the composite likelihood ratio (CLR) within each population was calculated in non-overlapping windows of 5kbp. We considered CLR peaks above the 95th percentile as putative selective sweeps. Signatures of selective sweeps can be located within a 10kb window centered on the true location of the sweep (Nielsen et al. 2005). Therefore, we considered an SV to be associated with a CLR peak if it overlaps or is within 5kb of a putative sweep window (Supplementary Fig. 7). We calculated Tajima’s D and nucleotide heterozygosity (π) for non-overlapping 5kbp windows using VCFtools v0.1.16 (Danecek et al. 2011). These summary statistics were calculated within each population and between samples with and without candidate SVs.

### Enrichment of SVs in Genomic Contexts

All SVs were classified into five mutually exclusive groups: exonic (fully contained within an exon), whole gene (overlaps at least one complete gene), partial gene (overlaps gene but not complete), intronic (fully contained within an intron), intergenic (fully contained within an intergenic region). To determine whether SVs are overrepresented or depleted in these genomic contexts, we compared the observed distribution to a null distribution generated by randomly shuffling the breakpoints of our SV call set 1000 times using BEDtools v2.30.0 *shuffle (Quinlan and Hall 2010)*.

### Enrichment of SVs near CLR peaks

To determine whether SVs segregating at allele frequencies greater than 25% are enriched near CLR peaks, we randomly shuffled the breakpoints of these SVs using BEDtools v2.30.0 *shuffle (Quinlan and Hall 2010)* and counted the number of SVs associated with a genomic window with CLR values higher than the 95th percentile. We repeated this 100,000 times for each population to generate null distributions. P-values were estimated using the following formula p=(r+1)/(n+1) (North et al. 2002), where r is the number of replicates in which the number of SVs associated with a CLR peak is greater than the observed number and n is the number of replicates.

### GO Term Enrichment Analysis

We performed GO enrichment analysis for genes affected by SVs segregating over 25% in a population and near CLR peaks using the topGO R package, v2.56.0. GO annotations for the UCI AnSteph reference genome were obtained from VectorBase.

### Inference of Time Since Onset of Selection

We applied an Approximate Bayesian computation (ABC) based approach adapted from Ormond et al. to estimate the time since the onset of selection (Ts) on the high-frequency esterase duplication (Ormond et al. 2016). While the duplication is high frequency in all populations, we used data from one mainland population, Trivandrum, for this calculation. We used one mainland population rather than all populations used in this study to minimize assumptions about the unknown demographic history of *An. stephensi* in India. Due to the single origin of the Esterase duplication allele, estimate of selective sweep from one population should be similar to estimates from other or all populations combined. First, we used *dadi (Gutenkunst et al. 2010)* to fit the demography model to the SFS of synonymous SNPs and estimate the model parameters. Based on the likelihood estimates of various demography models, we found that the best-fitting model was that of a population bottleneck followed by exponential growth. Using the estimate of effective population size of 1,757,732 inferred by *dadi*, we obtained an estimate for the strength of selection *s* from the inferred strength of selection by SweepFinder2. In particular, we used the following formula: s=r x ln(2Ne)/ɑ (Durrett and Schweinsberg 2004; Nielsen et al. 2005) where Ne is obtained from *dadi* estimated parameters, the intensity of selection ɑ of the CLR peak overlapping the duplication, and an assumed r=0.01cM/Mbp. To generate a prior distribution for Ts, 500,000 simulations were run with the coalescent simulation program MSMS (Ewing and Hermisson 2010), drawing Ts from a uniform distribution and incorporating the estimated s and demographic scenario inferred from *dadi*. Polymorphism data for the 20kb sequences flanking the esterase duplication were used for simulations. Summary statistics for simulations were calculated from MSMS output using the libsequence library (Thornton 2003) msstats function. Using the R package *pls* (Mevik and Wehrens 2007), a partial least squares method was applied to incorporate the most informative summary statistics into the ABC calculation. The posterior distribution for Ts was generated using an ABC rejection algorithm implemented by the R package *abc* (Csilléry et al. 2012). The point estimate for Ts was calculated from the mode of the posterior distribution.

## Data availability

All scripts and intermediate data files necessary to reproduce the work are available at https://github.com/chakrabortymlab/stephensi2024. The raw reads have been deposited to NCBI.

## Acknowledgments

We thank Rachel Moran and Susanta Ghosh for their helpful discussions. M.C. and A.S. were supported by funding from the National Institutes of Health grant R00GM129411 and start-up funds from Texas A&M to M.C. Mosquito collection fieldwork, and next-generation sequencing was funded by Tata Trusts to Tata Institute for Genetics and Society. We thank Sunita Swain and Sampath Kumar at the Tata Institute for Genetics and Society for their valuable suggestions.

## Author Contributions

M.C. and F.I. conceived the study. F.I. and N.K. contributed to the genomic data collection, sequencing, and variant calling. Y.L. performed the benchmarking analysis of SV detection methods. A.S. performed the population genetic analysis. A.S., M.C., and F. I. interpreted the results, and A.S. and M.C. wrote the manuscript. All authors reviewed and edited the manuscript.

## Notes

https://github.com/chakrabortymlab/stephensi2024

### Competing Interest Statement

The authors have declared no competing interest.

### Summary of Updates

The order of author names has been corrected.

## References

1. Abubakr M, Sami H, Mahdi I, Altahir O, Abdelbagi H, Mohamed NS, Ahmed A. 2022. The Phylodynamic and Spread of the Invasive Asian Malaria Vectors, Anopheles stephensi, in Sudan. Biology [Internet] 11. Available from: 10.3390/biology11030409

2. Abyzov A, Urban AE, Snyder M, Gerstein M. 2011. CNVnator: an approach to discover, genotype, and characterize typical and atypical CNVs from family and population genome sequencing. Genome Res. 21:974–984.

3. Ahmed W, Neelakanta G, Sultana H. 2021. Tetraspanins as Potential Therapeutic Candidates for Targeting Flaviviruses. Front. Immunol. 12:630571.

4. Alachiotis N, Pavlidis P. 2018. RAiSD detects positive selection based on multiple signatures of a selective sweep and SNP vectors. Communications Biology 1:1–11.

5. Anthony N, Unruh T, Ganser D, ffrench-Constant R. 1998. Duplication of the Rdl GABA receptor subunit gene in an insecticide-resistant aphid, Myzus persicae. Mol. Gen. Genet. 260:165–175.

6. Batra CP, Raghavendra K, Adak T, Singh OP, Singh SP, Mittal PK, Malhotra MS, Sharma RS, Subbarao SK. 2005. Evaluation of bifenthrin treated mosquito nets against anopheline and culicine mosquitoes. Indian J. Med. Res. 121:55–62.

7. Bohlen PJ, Scheu S, Hale CM, McLean MA, Migge S, Groffman PM, Parkinson D. 2004. Non-native invasive earthworms as agents of change in northern temperate forests. Front. Ecol. Environ. 2:427–435.

8. Bolger AM, Lohse M, Usadel B. 2014. Trimmomatic: a flexible trimmer for Illumina sequence data. Bioinformatics 30:2114–2120.

9. Bomblies K, Peichel CL. 2022. Genetics of adaptation. Proc. Natl. Acad. Sci. U. S. A. 119:e2122152119.

10. Bonizzoni M, Ochomo E, Dunn WA, Britton M, Afrane Y, Zhou G, Hartsel J, Lee M-C, Xu J, Githeko A, et al. 2015. RNA-seq analyses of changes in the Anopheles gambiae transcriptome associated with resistance to pyrethroids in Kenya: identification of candidate-resistance genes and candidate-resistance SNPs. Parasit. Vectors 8:474.

11. Carter TE, Yared S, Gebresilassie A, Bonnell V, Damodaran L, Lopez K, Ibrahim M, Mohammed S, Janies D. 2018. First detection of Anopheles stephensi Liston, 1901 (Diptera: culicidae) in Ethiopia using molecular and morphological approaches. Acta Trop. 188:180–186.

12. Chakraborty M, Emerson JJ, Macdonald SJ, Long AD. 2019. Structural variants exhibit widespread allelic heterogeneity and shape variation in complex traits. Nat. Commun. 10:4872.

13. Chakraborty M, Ramaiah A, Adolfi A, Halas P, Kaduskar B, Ngo LT, Jayaprasad S, Paul K, Whadgar S, Srinivasan S, et al. 2021. Hidden genomic features of an invasive malaria vector, Anopheles stephensi, revealed by a chromosome-level genome assembly. BMC Biol. 19:28.

14. Chakraborty M, VanKuren NW, Zhao R, Zhang X, Kalsow S, Emerson JJ. 2018. Hidden genetic variation shapes the structure of functional elements in Drosophila. Nat. Genet. 50:20–25.

15. Chen X, Schulz-Trieglaff O, Shaw R, Barnes B, Schlesinger F, Källberg M, Cox AJ, Kruglyak S, Saunders CT. 2016. Manta: rapid detection of structural variants and indels for germline and cancer sequencing applications. Bioinformatics 32:1220–1222.

16. Chung H, Sztal T, Pasricha S, Sridhar M, Batterham P, Daborn PJ. 2009. Characterization of *Drosophila melanogaster* cytochrome P450 genes. Proceedings of the National Academy of Sciences 106:5731–5736.

17. Cingolani P, Platts A, Wang LL, Coon M, Nguyen T, Wang L, Land SJ, Lu X, Ruden DM. 2012. A program for annotating and predicting the effects of single nucleotide polymorphisms, SnpEff: SNPs in the genome of Drosophila melanogaster strain w1118; iso-2; iso-3. Fly 6:80–92.

18. Coluzzi M. 1972. Inversion Polymorphism and Adult Emergence in *Anopheles stephensi*. Science 176:59–60.

19. Cridland JM, Macdonald SJ, Long AD, Thornton KR. 2013. Abundance and Distribution of Transposable Elements in Two Drosophila QTL Mapping Resources. Mol. Biol. Evol. 30:2311–2327.

20. Cridland JM, Thornton KR, Long AD. 2015. Gene expression variation in Drosophila melanogaster due to rare transposable element insertion alleles of large effect. Genetics 199:85–93.

21. Csilléry K, François O, Blum MGB. 2012. abc: an R package for approximate Bayesian computation (ABC). Methods Ecol. Evol. 3:475–479.

22. Daborn PJ, Yen JL, Bogwitz MR, Le Goff G, Feil E, Jeffers S, Tijet N, Perry T, Heckel D, Batterham P, et al. 2002. A single p450 allele associated with insecticide resistance in Drosophila. Science 297:2253–2256.

23. Danecek P, Auton A, Abecasis G, Albers CA, Banks E, DePristo MA, Handsaker RE, Lunter G, Marth GT, Sherry ST, et al. 2011. The variant call format and VCFtools. Bioinformatics 27:2156–2158.

24. Danecek P, Bonfield JK, Liddle J, Marshall J, Ohan V, Pollard MO, Whitwham A, Keane T, McCarthy SA, Davies RM, et al. 2021. Twelve years of SAMtools and BCFtools. Gigascience 10:giab008.

25. DeGiorgio M, Huber CD, Hubisz MJ, Hellmann I, Nielsen R. 2016. SweepFinder2: increased sensitivity, robustness and flexibility. Bioinformatics 32:1895–1897.

26. Derilus D, Impoinvil LM, Muturi EJ, McAllister J, Kenney J, Massey SE, Hemme R, Kothera L, Lenhart A. 2023. Comparative Transcriptomic Analysis of Insecticide-Resistant Aedes aegypti from Puerto Rico Reveals Insecticide-Specific Patterns of Gene Expression. Genes [Internet] 14. Available from: 10.3390/genes14081626

27. Durrett R, Schweinsberg J. 2004. Approximating selective sweeps. Theor. Popul. Biol. 66:129–138.

28. Dykes CL, Sharma G, Behera AK, Kapoor N, Paine MJI, Donnelly MJ, Singh OP. 2022. Tandem duplication of a genomic region encoding glutathione S-transferase epsilon-2 and-4 genes in DDT-resistant Anopheles stephensi strain from India. Sci. Rep. 12:1–14.

29. Emerson JJ, Cardoso-Moreira M, Borevitz JO, Long M. 2008. Natural Selection Shapes Genome-Wide Patterns of Copy-Number Polymorphism in *Drosophila melanogaster*. Science 320:1629–1631.

30. Enayati A, Hanafi-Bojd AA, Sedaghat MM, Zaim M, Hemingway J. 2020. Evolution of insecticide resistance and its mechanisms in Anopheles stephensi in the WHO Eastern Mediterranean Region. Malar. J. 19:258.

31. Ewing G, Hermisson J. 2010. MSMS: a coalescent simulation program including recombination, demographic structure and selection at a single locus. Bioinformatics 26:2064–2065.

32. Ganesh KN, Vijayan VA, Urmila J, Gopalan N, Prakash S. 2002. Role of esterases and monooxygenase in the deltamethrin resistance in Anopheles stephensi Giles (1908), at Mysore. Indian J. Exp. Biol. 40:583–588.

33. Garrison E, Marth G. 2012. Haplotype-based variant detection from short-read sequencing. arXiv [q-bio.GN] [Internet]. Available from: http://arxiv.org/abs/1207.3907

34. Dharmasiri AG, Perera AY, Harishchandra J, Herath H, Aravindan K, Jayasooriya HTR, Ranawaka GR, Hewavitharane M. 2017. First record of Anopheles stephensi in Sri Lanka: a potential challenge for prevention of malaria reintroduction. Malar. J. 16:326.

35. Gunasekaran K, Jambulingam P, Srinivasan R, Sadanandane C, Doss PSB, Sabesan S, Balaraman K, Das PK. 2005. Malaria receptivity in the tsunami-hit coastal villages of southern India. Lancet Infect. Dis. 5:531–532.

36. Gutenkunst R, Hernandez R, Williamson S, Bustamante C. 2010. Diffusion Approximations for Demographic Inference: DaDi. Nature Precedings:1–1.

37. Habtewold T, Wagah M, Tambwe MM, Moore S, Windbichler N, Christophides G, Johnson H, Heaton H, Collins J, Krasheninnikova K, et al. 2023. A chromosomal reference genome sequence for the malaria mosquito, Anopheles gambiae, Giles, 1902, Ifakara strain. Wellcome Open Res 8:74.

38. Hardstone MC, Komagata O, Kasai S, Tomita T, Scott JG. 2010. Use of isogenic strains indicates CYP9M10 is linked to permethrin resistance in Culex pipiens quinquefasciatus. Insect Mol. Biol. 19:717–726.

39. Harringmeyer OS, Hoekstra HE. 2022. Chromosomal inversion polymorphisms shape the genomic landscape of deer mice. Nat Ecol Evol 6:1965–1979.

40. Hayes KA, Joshi RC, Thiengo SC, Cowie RH. 2008. Out of South America: multiple origins of non-native apple snails in Asia. Divers. Distrib. 14:701–712.

41. Heilig M, Sturiale SL, Marzec S, Holzapfel CM, Bradshaw WE, Meuti ME, Armbruster PA. 2024. Phenotypic variation in biting behavior associated with differences in expression of olfactory genes in the vector mosquito, Aedes albopictus (Diptera: Culicidae). J. Med. Entomol. 61:367–376.

42. Hemingway J, Karunaratne SH. 1998. Mosquito carboxylesterases: a review of the molecular biology and biochemistry of a major insecticide resistance mechanism. Med. Vet. Entomol. 12:1–12.

43. Hemingway J, Ranson H. 2000. Insecticide resistance in insect vectors of human disease. Annu. Rev. Entomol. 45:371–391.

44. Hemming-Schroeder E, Ahmed A. 2023. Anopheles stephensi in Africa: vector control opportunities for cobreeding An. stephensi and Aedes arbovirus vectors. Trends Parasitol. 39:86–90.

45. Hoskins RA, Carlson JW, Wan KH, Park S, Mendez I, Galle SE, Booth BW, Pfeiffer BD, George RA, Svirskas R, et al. 2015. The Release 6 reference sequence of the Drosophila melanogaster genome. Genome Res. 25:445–458.

46. Huber CD, Kim BY, Marsden CD, Lohmueller KE. 2017. Determining the factors driving selective effects of new nonsynonymous mutations. Proc. Natl. Acad. Sci. U. S. A. 114:4465–4470.

47. Ishtiaq F, Swain S, Kumar SS. 2021. Anopheles stephensi (Asian Malaria Mosquito). Trends Parasitol. 37:571–572.

48. Kent WJ, Baertsch R, Hinrichs A, Miller W, Haussler D. 2003. Evolution’s cauldron: Duplication, deletion, and rearrangement in the mouse and human genomes. Proceedings of the National Academy of Sciences 100:11484–11489.

49. Kirsche M, Prabhu G, Sherman R, Ni B, Battle A, Aganezov S, Schatz MC. 2023. Jasmine and Iris: population-scale structural variant comparison and analysis. Nat. Methods 20:408–417.

50. Kolaczinski J, Al-Eryani S, Chanda E, Fernandez-Montoya L. 2021. Comment on: Emergence of the invasive malaria vector Anopheles stephensi in Khartoum State, Central Sudan. Parasit. Vectors 14:588.

51. Labbé P, Berthomieu A, Berticat C, Alout H, Raymond M, Lenormand T, Weill M. 2007. Independent duplications of the acetylcholinesterase gene conferring insecticide resistance in the mosquito Culex pipiens. Mol. Biol. Evol. 24:1056–1067.

52. Layer RM, Chiang C, Quinlan AR, Hall IM. 2014. LUMPY: a probabilistic framework for structural variant discovery. Genome Biol. 15:R84.

53. Lee CE. 2002. Evolutionary genetics of invasive species. Trends Ecol. Evol. 17:386–391.

54. Lee CE, Kiergaard M, Gelembiuk GW, Eads BD, Posavi M. 2011. Pumping ions: rapid parallel evolution of ionic regulation following habitat invasions. Evolution 65:2229–2244.

55. Li H. 2013. Aligning sequence reads, clone sequences and assembly contigs with BWA-MEM. arXiv [q-bio.GN] [Internet]. Available from: http://arxiv.org/abs/1303.3997

56. Liu N. 2015. Insecticide Resistance in Mosquitoes: Impact, Mechanisms, and Research Directions. Annu. Rev. Entomol. 60:537–559.

57. Mahmood F, Sakai RK. 1984. Inversion polymorphisms in natural populations of Anopheles stephensi. Can. J. Genet. Cytol. 26:538–546.

58. Mevik B-H, Wehrens R. 2007. The pls package: Principal component and partial least squares regression in R. J. Stat. Softw. 18:1–23.

59. Morrison LW, Porter SD, Daniels E, Korzukhin MD. 2004. Potential Global Range Expansion of the Invasive Fire Ant, Solenopsis invicta. Biol. Invasions 6:183–191.

60. Newcomb RD, Gleeson DM, Yong CG, Russell RJ, Oakeshott JG. 2005. Multiple mutations and gene duplications conferring organophosphorus insecticide resistance have been selected at the Rop-1 locus of the sheep blowfly, Lucilia cuprina. J. Mol. Evol. 60:207–220.

61. Nielsen R, Williamson S, Kim Y, Hubisz MJ, Clark AG, Bustamante C. 2005. Genomic scans for selective sweeps using SNP data. Genome Res. 15:1566–1575.

62. North BV, Curtis D, Sham PC. 2002. A note on the calculation of empirical P values from Monte Carlo procedures. Am. J. Hum. Genet. 71:439–441.

63. Ormond L, Foll M, Ewing GB, Pfeifer SP, Jensen JD. 2016. Inferring the age of a fixed beneficial allele. Mol. Ecol. 25:157–169.

64. Park SG, Hannenhalli S, Choi SS. 2014. Conservation in first introns is positively associated with the number of exons within genes and the presence of regulatory epigenetic signals. BMC Genomics 15:526.

65. Prasad KM, Raghavendra K, Verma V, Velamuri PS, Pande V. 2017. Esterases are responsible for malathion resistance in Anopheles stephensi: A proof using biochemical and insecticide inhibition studies. J. Vector Borne Dis. 54:226–232.

66. Prentis PJ, Wilson JRU, Dormontt EE, Richardson DM, Lowe AJ. 2008. Adaptive evolution in invasive species. Trends Plant Sci. 13:288–294.

67. Quinlan AR, Hall IM. 2010. BEDTools: a flexible suite of utilities for comparing genomic features. Bioinformatics 26:841–842.

68. Raghavendra K, Velamuri PS, Verma V, Elamathi N, Barik TK, Bhatt RM, Dash AP. 2017. Temporo-spatial distribution of insecticide-resistance in Indian malaria vectors in the last quarter-century: Need for regular resistance monitoring and management. J. Vector Borne Dis. 54:111.

69. Rahi M, Sharma A. 2022. Malaria control initiatives that have the potential to be gamechangers in India’s quest for malaria elimination. Lancet Reg Health Southeast Asia 2:100009.

70. Ramasamy R, Thiruchenthooran V, Jayadas TTP, Eswaramohan T, Santhirasegaram S, Sivabalakrishnan K, Naguleswaran A, Uzest M, Cayrol B, Voisin SN, et al. 2021. Transcriptomic, proteomic and ultrastructural studies on salinity-tolerant Aedes aegypti in the context of rising sea levels and arboviral disease epidemiology. BMC Genomics 22:253.

71. Rausch T, Zichner T, Schlattl A, Stütz AM, Benes V, Korbel JO. 2012. DELLY: structural variant discovery by integrated paired-end and split-read analysis. Bioinformatics 28:i333–i339.

72. Reznick DN, Ghalambor CK. 2001. The population ecology of contemporary adaptations: what empirical studies reveal about the conditions that promote adaptive evolution. Genetica 112:183–198.

73. Schmidt JM, Good RT, Appleton B, Sherrard J, Raymant GC, Bogwitz MR, Martin J, Daborn PJ, Goddard ME, Batterham P, et al. 2010. Copy number variation and transposable elements feature in recent, ongoing adaptation at the Cyp6g1 locus. PLoS Genet. 6:e1000998.

74. Schwartz S, Kent WJ, Smit A, Zhang Z, Baertsch R, Hardison RC, Haussler D, Miller W. 2003. Human-mouse alignments with BLASTZ. Genome Res. 13:103–107.

75. Sharma S, Hamzakoya KK. 2001. Geographical spread of Anopheles stephensi, vector of urban malaria, and Aedes aegypti, vector of dengue/DHF, in the Arabian sea islands of Lakshadweep, India. Available from: https://apps.who.int/iris/bitstream/handle/10665/204930/B0223.pdf#page=95

76. Shukla HG, Chakraborty M, Emerson J. 2024. Genetic variation in recalcitrant repetitive regions of the Drosophila melanogaster genome. bioRxiv [Internet]. Available from: http://biorxiv.org/lookup/doi/10.1101/2024.06.11.598575

77. Sinka ME, Pironon S, Massey NC, Longbottom J, Hemingway J, Moyes CL, Willis KJ. 2020. A new malaria vector in Africa: Predicting the expansion range of Anopheles stephensi and identifying the urban populations at risk. Proceedings of the National Academy of Sciences [Internet]:202003976. Available from: 10.1073/pnas.2003976117

78. Sivabalakrishnan K, Thanihaichelvan M, Tharsan A, Eswaramohan T, Ravirajan P, Hemphill A, Ramasamy R, Surendran SN. 2023. Resistance to the larvicide temephos and altered egg and larval surfaces characterize salinity-tolerant Aedes aegypti. Sci. Rep. 13:1–12.

79. Slater GSC, Birney E. 2005. Automated generation of heuristics for biological sequence comparison. BMC Bioinformatics 6:31.

80. Smith KE, VanEkeris LA, Okech BA, Harvey WR, Linser PJ. 2008. Larval anopheline mosquito recta exhibit a dramatic change in localization patterns of ion transport proteins in response to shifting salinity: a comparison between anopheline and culicine larvae. J. Exp. Biol. 211:3067–3076.

81. Stephan W. 2019. Selective Sweeps. Genetics 211:5–13.

82. Surendran SN, Sivabalakrishnan K, Sivasingham A, Jayadas TTP, Karvannan K, Santhirasegaram S, Gajapathy K, Senthilnanthanan M, Karunaratne SP, Ramasamy R. 2019. Anthropogenic Factors Driving Recent Range Expansion of the Malaria Vector Anopheles stephensi. Front Public Health 7:53.

83. Takken W, Lindsay S. 2019. Increased Threat of Urban Malaria from Anopheles stephensi Mosquitoes, Africa. Emerg. Infect. Dis. 25:1431–1433.

84. Thakare A, Ghosh C, Alalamath T, Kumar N, Narang H, Whadgar S, Paul K, Shrotri S, Kumar S, Soumya M, et al. 2022. The genome trilogy of Anopheles stephensi, an urban malaria vector, reveals structure of a locus associated with adaptation to environmental heterogeneity. Sci. Rep. 12:1–16.

85. Thornton K. 2003. Libsequence: a C++ class library for evolutionary genetic analysis. Bioinformatics 19:2325–2327.

86. Thorvaldsdóttir H, Robinson JT, Mesirov JP. 2013. Integrative Genomics Viewer (IGV): high-performance genomics data visualization and exploration. Brief. Bioinform. 14:178–192.

87. Tiwari S, Ghosh SK, Ojha VP, Dash AP, Raghavendra K. 2010. Reduced susceptibility to selected synthetic pyrethroids in urban malaria vector Anopheles stephensi: a case study in Mangalore city, South India. Malar. J. 9:179.

88. Van’t Hof AE, Campagne P, Rigden DJ, Yung CJ, Lingley J, Quail MA, Hall N, Darby AC, Saccheri IJ. 2016. The industrial melanism mutation in British peppered moths is a transposable element. Nature 534:102–105.

89. Weetman D, Mitchell SN, Wilding CS, Birks DP, Yawson AE, Essandoh J, Mawejje HD, Djogbenou LS, Steen K, Rippon EJ, et al. 2015. Contemporary evolution of resistance at the major insecticide target site gene Ace-1 by mutation and copy number variation in the malaria mosquito Anopheles gambiae. Mol. Ecol. 24:2656– 2672.

90. Wilding CS, Smith I, Lynd A, Yawson AE, Weetman D, Paine MJI, Donnelly MJ. 2012. A cis-regulatory sequence driving metabolic insecticide resistance in mosquitoes: functional characterisation and signatures of selection. Insect Biochem. Mol. Biol. 42:699–707.

91. Yared S, Gebressielasie A, Damodaran L, Bonnell V, Lopez K, Janies D, Carter TE. 2020. Insecticide resistance in Anopheles stephensi in Somali Region, eastern Ethiopia. Malar. J. 19:180.

92. Zimmer CT, Garrood WT, Singh KS, Randall E, Lueke B, Gutbrod O, Matthiesen S, Kohler M, Nauen R, Emyr Davies TG, et al. 2018. Neofunctionalization of Duplicated P450 Genes Drives the Evolution of Insecticide Resistance in the Brown Planthopper. Curr. Biol. 28:268.

93. Alexa A, Rahnenfuhrer J. 2024. topGO: Enrichment Analysis for Gene Ontology. R package version 2.56.0.

