## Supplementary Information for "Genome structural variants shape adaptive success of an invasive urban malaria vector *Anopheles stephensi*"

### Supplementary Figures

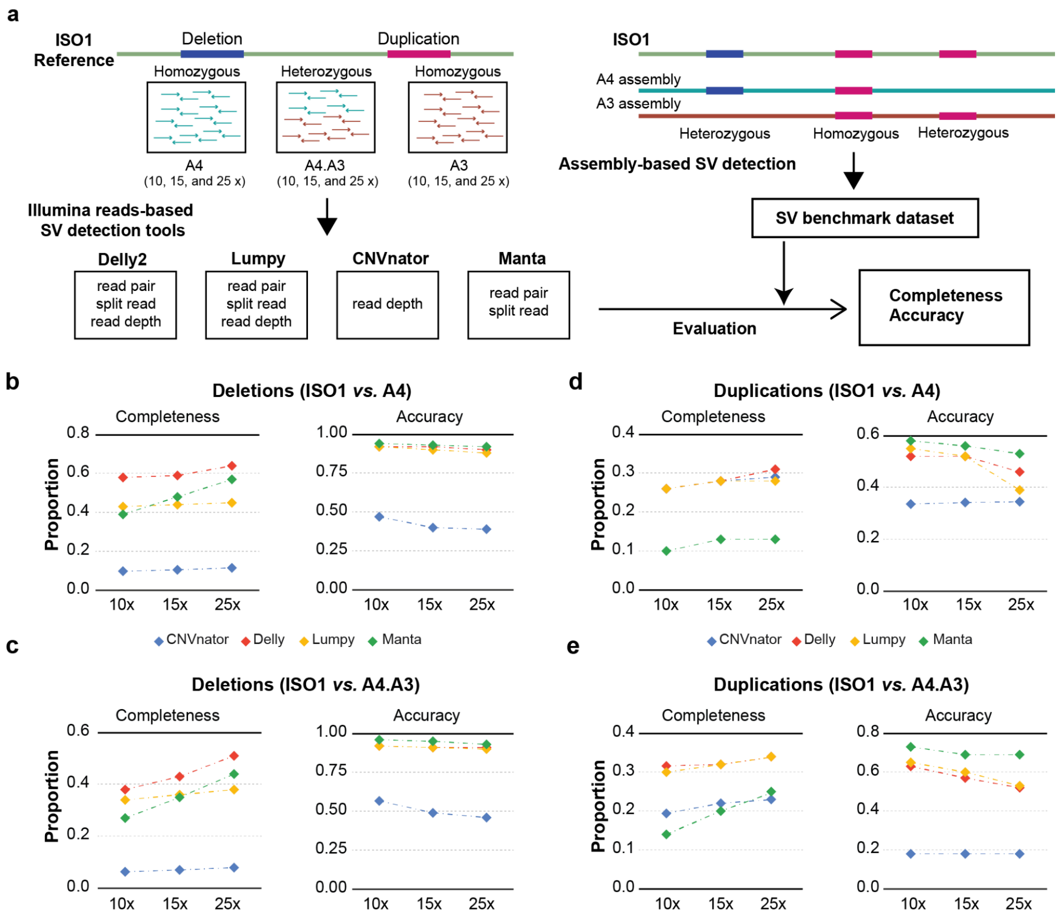


**Supplementary Figure 1.** Strategy and workflow of the benchmarking analysis. **a.** Paired-end Illumina reads from two inbred *Drosophila melanogaster* lines were sampled at three different coverage depths. Heterozygous data was simulated by equally combining reads from two lines. The performance of 4 short-read mapping SV callers was evaluated by comparing them to SV calls from assembly-based SV detection. Completeness refers to the proportion of total SVs in the assembly discoverable by the short-read caller. Accuracy refers to the proportion of SVs detected by the short reads also present in the assembly. **b.** Evaluation of deletion calls from A4 reads mapped to the ISO1 reference assembly. **c.** Evaluation of deletion calls from simulated A3/A4 heterozygous reads mapped to the ISO1 reference assembly. **d.** Evaluation of duplication calls from A4 reads mapped to the ISO1 reference assembly. **e.** Evaluation of duplication calls from simulated A3/A4 heterozygous reads mapped to the ISO1 reference assembly.

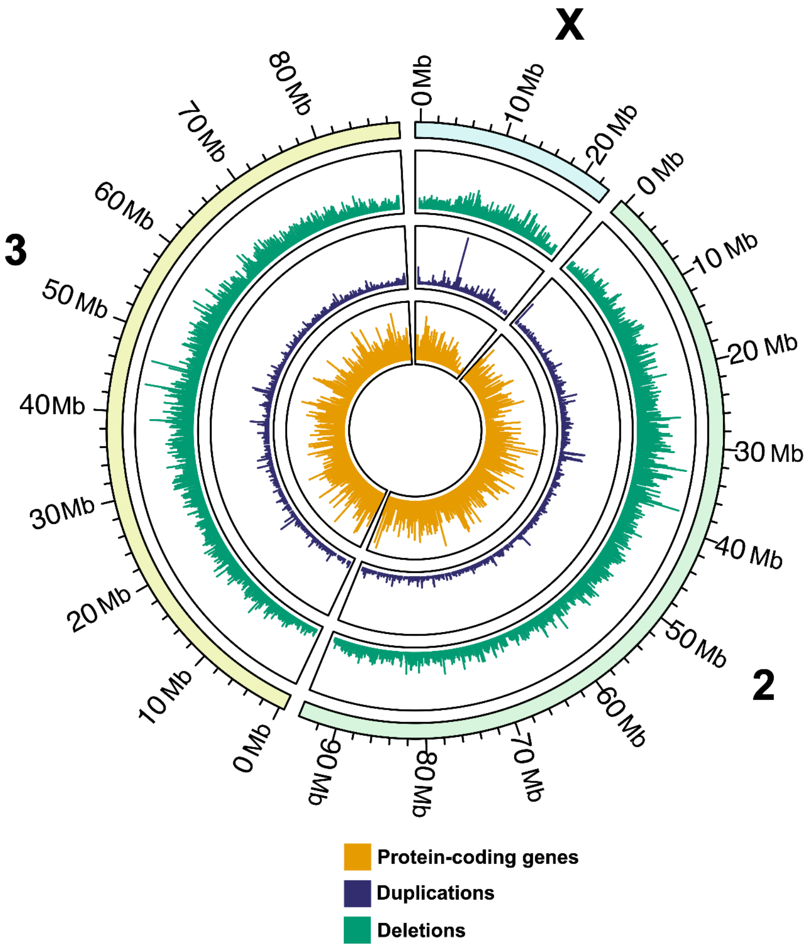


**Supplementary Figure 2.** Distribution of CNVs (relative to the AnStephUCI reference) of length 100bp-100Kbp and genes across chromosomes 2, 3, and X. Tracks show the number of deletions (green), duplications (purple), and protein-coding genes (orange) per 100 Kbp window in the genome.

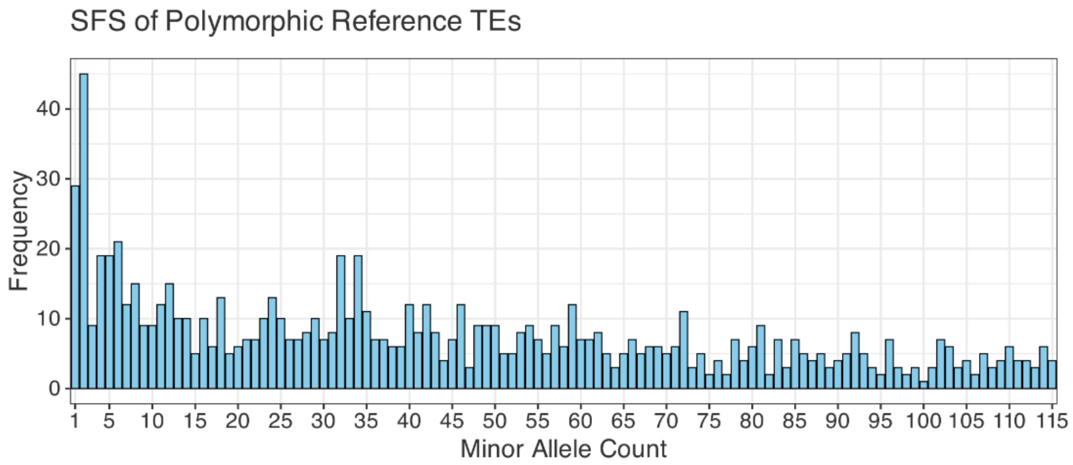

**Supplementary Figure 3.** Histogram of minor allele counts for polymorphic transposable element (TE) insertions present in the AnStephUCI reference genome.

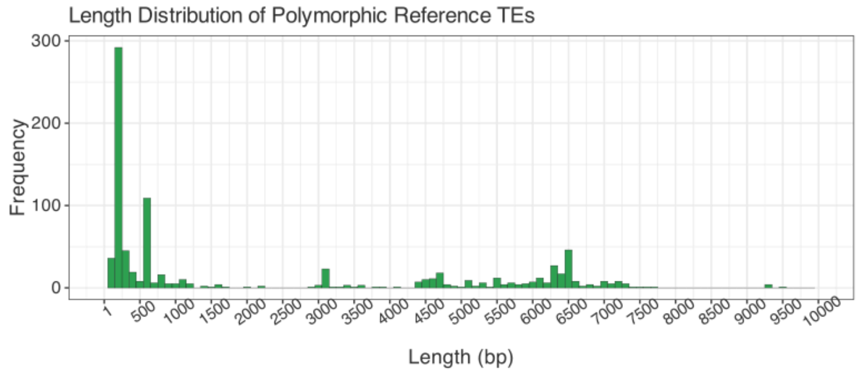


**Supplementary Figure 4.** Binned histogram of polymorphic reference TE lengths under 10 kb. The peaks near 3 kb, 4.5 kb, and 6.5 kb likely represent lengths of major active full-length TEs in *An. stephensi*.

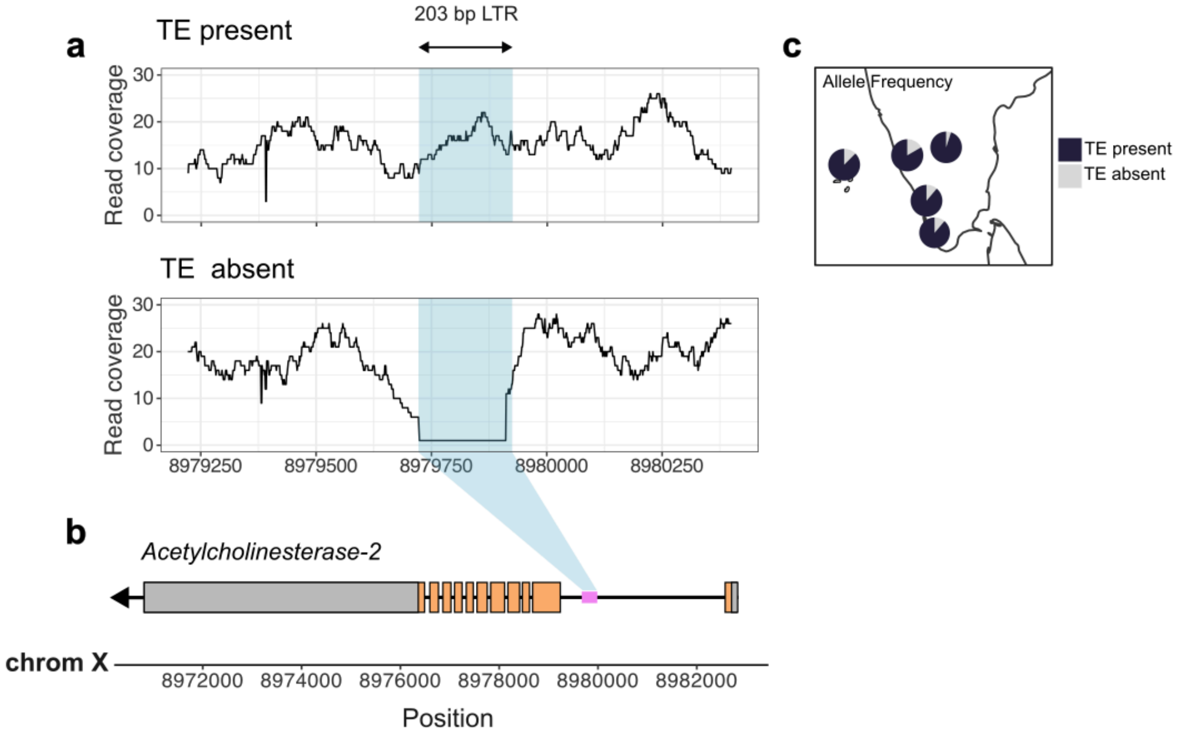


**Supplementary Figure 5. a.** Read coverage for a polymorphic TE insertion in the *Acetylcholinesterase-2* gene. The TE sequence is present in the UCI reference genome. Therefore, the absence of the TE insertion appears as a deletion that completely overlaps the TE fragment. **b.** This LTR retrotransposon fragment is inserted into the first intron of the gene. The length of the first intron suggests the presence of regulatory sites, which could potentially be affected by the LTR fragment. **c.** The allele frequency of the TE fragment ( 1- deletion frequency) in the population samples.

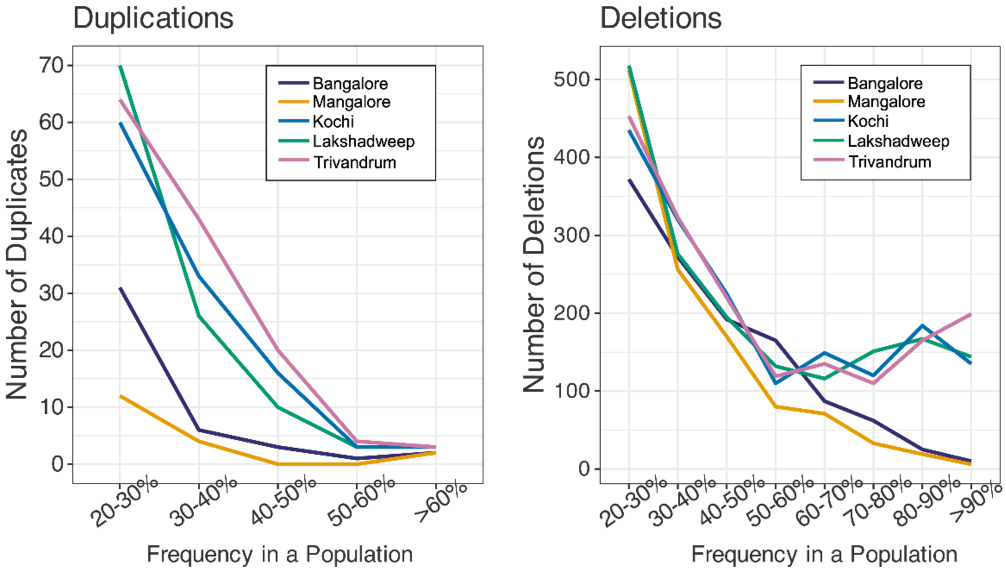


**Supplementary Figure 6.** Number of SVs segregating at frequencies greater than 20% in each population. The elevated number of deletions segregating at high frequencies in Kochi, Lakshadweep, and Trivandrum may include low-frequency insertions in the reference genome.

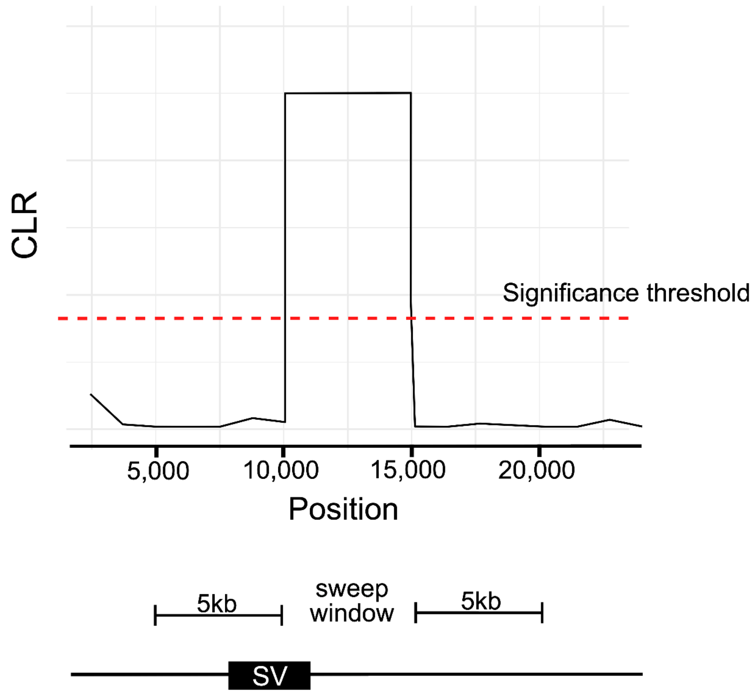


**Supplementary Figure 7.** Schematic demonstrating how it was determined whether an SV is associated with a CLR peak. The actual location of a sweep can be ~10 kbp from the CLR peak identified by SweepFinder2. Therefore, we included 5kbp on either side of the sweep window and overlapped it with our SV call set.

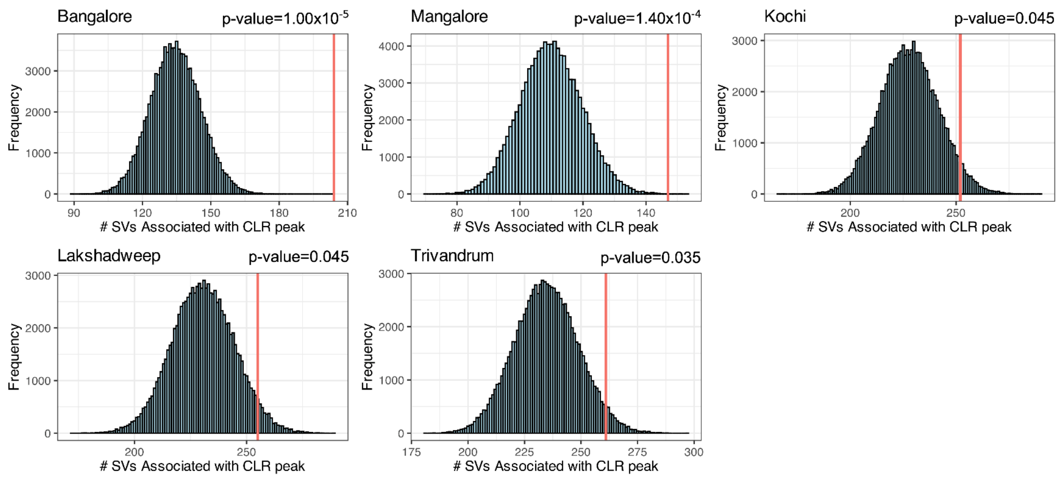


**Supplementary Figure 8.** Coordinates of SVs over 25% allele frequency in a population were shuffled 100,000 times. The number of SVs associated with CLR peaks was counted per run to generate null distributions for each population. Red lines represent the observed number of SVs over 25% allele frequency associated with CLR peaks.

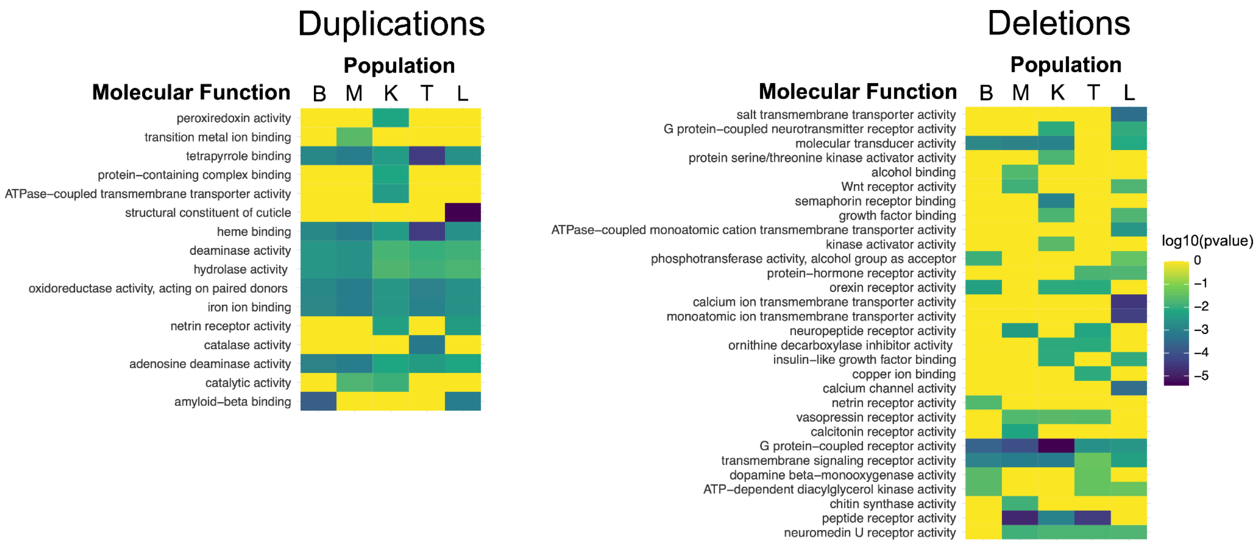


**Supplementary Figure 9.** GO term enrichment analysis for genes overlapped by duplications with allele frequency over 25% and associated with CLR peaks.

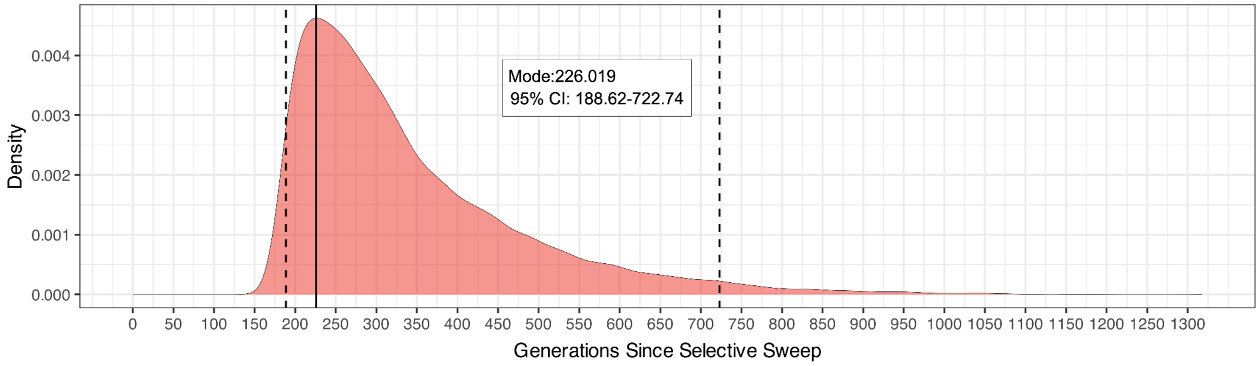


**Supplementary Figure 10.** The posterior probability distribution for the estimated age of a selective sweep in the Trivandrum population. This sweep is associated with a high-frequency duplication of carboxylesterases. The mode of the distribution is marked by the solid line, and 95% confidence intervals are marked by dotted lines.

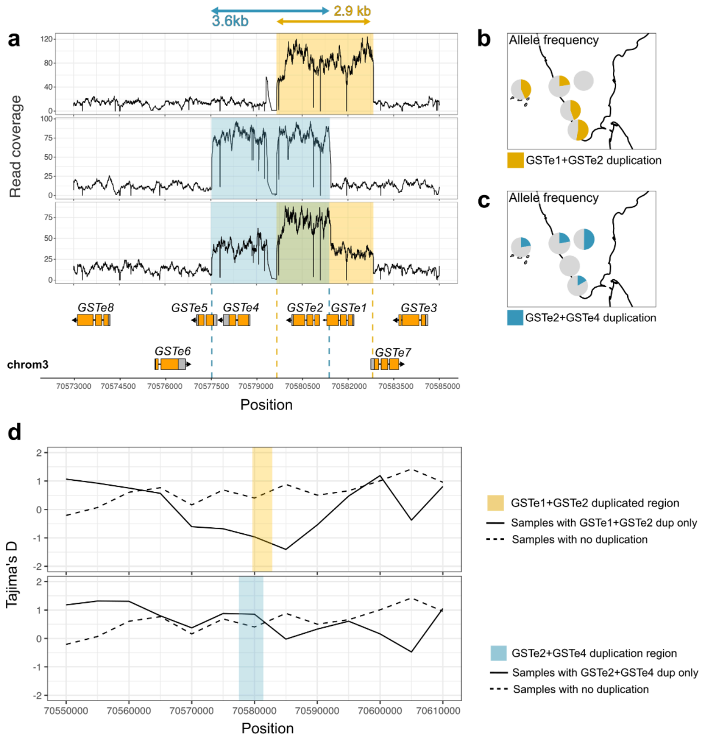


**Supplementary Figure 11. a**. Duplication CNVs in an array of epsilon Glutathione-S Transferase (GSTe) genes. b. One duplication (top) copies a sequence containing two full-length GSTes and partially overlaps another duplication allele (middle). 21 samples appear to have both alleles or a recombinant allele (bottom). **b.** Allele frequency of the duplication, which copies *GSTe1* and *GSTe2*. **c.** Allele frequency of the duplication copies *GSTe2, GSTe4,* and partial sequences. **d.** Reduced Tajima's D flanking the *GSTe1+GSTe2* duplication is indicative of positive selection.

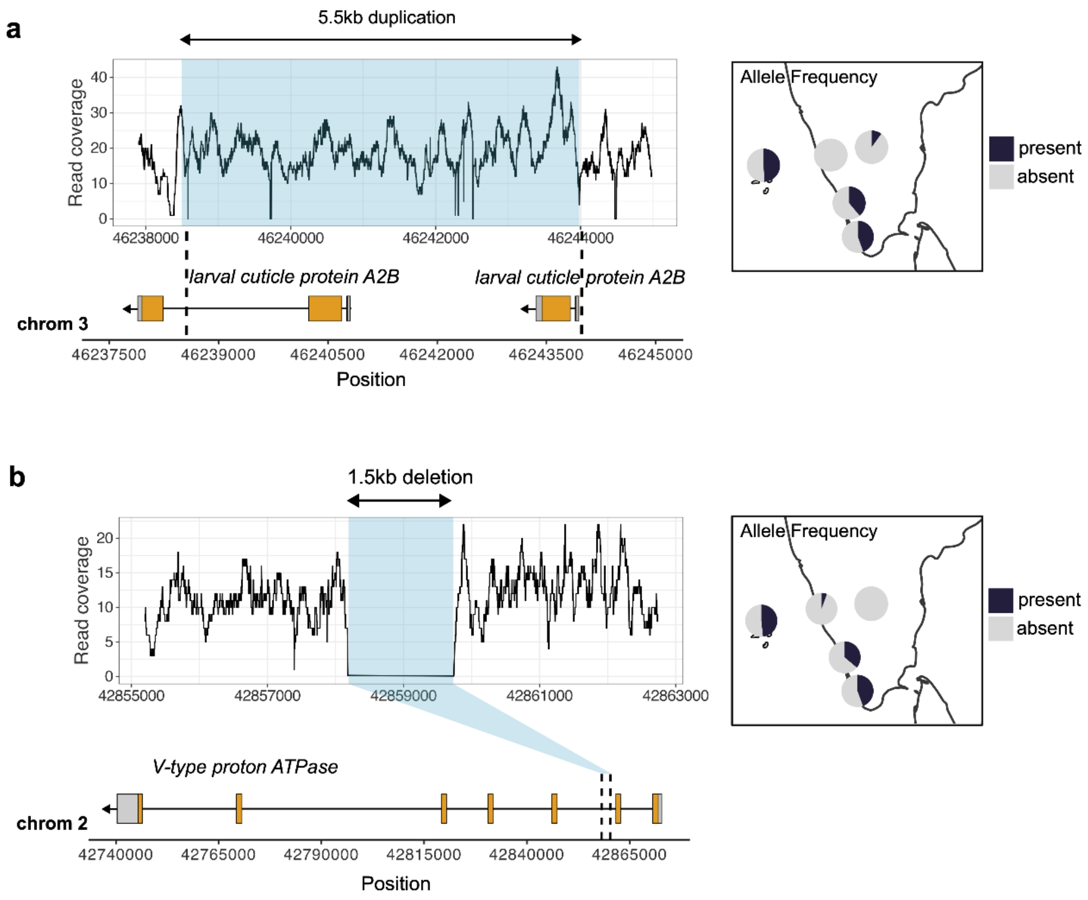


**Supplementary Figure 12.** Two candidate SVs found in coastal mainland and island populations. A duplication overlapping larval cuticle proteins (**a**) and an intronic deletion in an enzyme involved in osmoregulation (**b**) segregate at high frequencies in Kochi, Trivandrum, and Lakshadweep but are absent or low frequency in more inland populations.

### Supplementary Tables

**Supplementary Table 1.** Sample IDs and coverage.

**Supplementary Table 2.** IndCh SVs validated using PacBio long read and 20x Illumina short read coverage.

**Supplementary Table 3.** Duplication CNV coordinates, allele frequency per population, associated CLR if above the 95 percentile, and gene overlap.

**Supplementary Table 4.** Deletion CNV coordinates, allele frequency per population, associated CLR if above the 95 percentile, and gene overlap.

**Supplementary Table 5.** Polymorphic reference TE coordinates, allele frequency per population (1 – deletion frequency), associated CLR if above the 95 percentile, and geneoverlap.

**Supplementary Table 6.** Missense SNP positions, allele frequency per population, associated CLR if above the 95 percentile, and gene overlap.

**Supplementary Table 7**. Key for main figure 3C, esterase duplication gene tree.

| **Full ID** | **Short ID** |
|---|---|
| KIL0314D.S40 | L1 |
| KIL0314C.S32 | L2 |
| KIL0315D.S44 | L3 |
| KIL0340A.S45 | L4 |
| KIL0315A.S41 | L5 |
| KIL0314A.S30 | L6 |
| KIL0315B.S42 | L7 |
| C82.31.S69 | K1 |
| UCI.reference | REF |
| AGT0091A.S19 | L8 |
| AGT0091E.S23 | L9 |
| AGT0013B.S16 | L10 |
| AGT0013A.S15 | L11 |
| MCY0093A.S8 | L12 |
| AGT0091C.S21 | L13 |
| AGT0091D.S22 | L14 |
| AGT0091B.S20 | L15 |
| CHT0007J.S28 | L16 |
| CHT0007I.S27 | L17 |
| CHT0007G.S39 | L18 |
| CHT0007F.S38 | L19 |
| CHT0007K.S29 | L20 |
| CHT0007D.S36 | L21 |
| KVT0211.S6 | L22 |
| KVT0268A.S8 | L23 |
| AND0289.S4 | L24 |
| CHT0007C.S35 | L25 |
| KVT0158.S3 | L26 |

| | |
|---|---|
| KLP0003A.S46 | L27 |
| CHT0007H.S26 | L28 |
| KLP0003C.S48 | L29 |
| TVM7AE1.S9 | T1 |
| TVM25AE1.S11 | T2 |
| TVM111.S93 | T3 |
| TVM106.S90 | T4 |
| TVM105.S89 | T5 |
| TVM103.S87 | T6 |
| TVM109.S91 | T7 |
| TVM112.S94 | T8 |
| TVM110.S92 | T9 |
| TVM085.S84 | T10 |
| TVM104.S88 | T11 |
| TVM21AE1.S10 | T12 |
| TVM102.S86 | T13 |
| M10 | M10 |
| TVM086 | T14 |
| M6 | M6 |
| AND0301A.S6 | L30 |
| AND0089.S3 | L31 |
| AND0211.S5 | L32 |
| AND0046A.S1 | L33 |
| C82.63.S72 | K2 |
| C82.18.S68 | K3 |
| C82.7.S67 | K4 |
| C18.3.S95 | K5 |
| C82.64.S73 | K6 |
| C67.4.S77 | K7 |
| C66.27.S66 | K8 |
| C82.37.S70 | K9 |
| C66.19.S64 | K10 |
| M9 | M9 |
| M8 | M8 |
| M5 | M5 |
| M1 | M1 |
| B4 | B4 |
| B10 | B10 |
| B6 | B6 |

| | |
|---|---|
| B9 | B9 |
| KVT0157.S2 | L34 |
| KVT.O185.S3 | L35 |
| KVT0186.S4 | L36 |
| CHT0007A.S33 | L37 |
| KVT0156.S1 | L38 |
| BTR002.S80 | L39 |
